# Pro- and anti-inflammatory macrophages adjust UCP2 protein levels based on their intrinsic metabolism and available metabolites

**DOI:** 10.1101/2025.09.08.674165

**Authors:** Jila Nasirzade, Felix Sternberg, Andrea Vogel, Roko Sango, Taraneh Beikbaghban, Thomas Kolbe, Thomas Rattei, Thomas Weichhart, Elena E. Pohl

## Abstract

The immune and metabolic responses of macrophages are closely linked, and mitochondria play a key role in polarizing them into pro-inflammatory (classical) and anti-inflammatory (alternative) states. Mitochondrial uncoupling protein 2 (UCP2) is involved in regulating macrophage inflammation and glucose metabolism; however, its regulatory mechanisms are unclear. We found that inflammatory stimuli reduce UCP2 expression and oxygen consumption rates (OCR), indicating mitochondrial suppression. Conversely, IL-4-activated macrophages displayed higher UCP2 levels and enhanced respiration. Under glucose deprivation, LPS-stimulated macrophages retained mitochondrial activity despite lower UCP2 levels. Pyruvate emerged as a key regulator of UCP2, blocking its mitochondrial entry reduced UCP2 expression. Additionally, hypoxia markedly decreased UCP2 levels in IL-4-activated macrophages, suggesting that hypoxia contributes to UCP2 suppression in pro-inflammatory macrophages. Notably, pro-inflammatory macrophages exhibit reduced reliance on UCP2 due to suppressed mitochondrial respiration. Pyruvate regulates UCP2 expression, highlighting the connection between glycolysis and mitochondrial metabolism. These findings may inform therapeutic strategies for diseases involving immune dysregulation.

## 1. Introduction

Immune responses and metabolic regulation are intricately linked, both essential for the proper functioning of the organism. Disruptions in these processes can lead to a range of immunometabolism-related diseases, including cancer, obesity and diabetes. Immune cells adapt to dietary changes by modifying lipid and/or glucose metabolism, which, in turn, affects the intrinsic metabolism of surrounding cells crucial for maintaining metabolic homeostasis (Hotamisligil, 2017; Seufert et al., 2022). Numerous studies have underscored the role of macrophages in the glucose metabolism of obese mice (Choi et al., 2022; Li et al., 2023; Weisberg et al., 2003; Xu et al., 2003).

The immune and metabolic responses in macrophages are tightly regulated by mitochondria. In vitro, lipopolysaccharide (LPS) and interferon γ (INFγ) are used to polarize macrophages to a pro-inflammatory phenotype (LPS-MΦs), resulting in a shift from glucose-dependent oxidative phosphorylation (OxPhos) to aerobic glycolysis, a process known as the Warburg effect (Kelly and O’Neill, 2015). This phenotype has an impaired TCA cycle (Jha et al., 2015), making it more dependent on ATP produced by glycolysis rather than OxPhos in mitochondria (Newsholme, 2021). On the other hand, anti-inflammatory macrophages polarized in vitro by IL4 and IL13 stimulation (IL4-MΦs), are highly dependent on OxPhos and use glutamine as a major source of their metabolism (Jha et al., 2015). Macrophages can rapidly switch from a catabolic to anabolic state to support the immune cell phenotype, and mitochondria help in their polarization and adaptation to environmental changes (Lundahl et al., 2022). Although distinct metabolic profiles among macrophage subtypes are increasingly recognized, the mechanisms governing their selective metabolite utilization remain poorly understood.

Uncoupling protein 2 (UCP2) is a member of the large family of mitochondrial anion carriers (SLC25) and is located in the inner membrane of mitochondria. Although UCP2 was first discovered in the late 1990s (Fleury et al., 1997), its biological function remains a subject of ongoing debate. UCP2 has been shown to be abundant in cells that primarily rely on glycolysis, have high proliferation rates, and possess stemness characteristics (Hilse et al., 2020; Pohl et al., 2019), such as cancer cells (Beikbaghban et al., 2024; Esteves et al., 2015; Horimoto et al., 2004), stem cells (Rupprecht et al., 2014; Yu et al., 2013; Zhang et al., 2011) and immune cells, including monocytes, B cells, T cells and microglia (Maes et al., 2023; Rupprecht et al., 2012). Additionally, repeated stimulation has been shown to increase UCP2 abundance in T-cells (Rupprecht et al., 2012; Rupprecht et al., 2014).

UCP2 knockout experiments indicated the role of UCP2 in immune response, as UCP2 knockout mice were observed to exhibit increased resistance to *Toxoplasma gondii* infection (Arsenijevic et al., 2000). These knockout mice also showed decreased level of inflammatory cytokines such as IL1β and IL6 (Emre et al., 2007a; Emre et al., 2007b; van Dierendonck et al., 2020; Yan et al., 2020). Further evidence supporting the role of UCP2 in macrophage immune responses comes from studies showing an increase in UCP2 expression following LPS injection (Couplan et al., 2002; Pecqueur et al., 2001). Recent study has shown that microglia-specific UCP2 knockout mice exhibit significant mitochondrial hyperfusion following optical nerve crush (Maes et al., 2023). This effect was linked to an increased reliance on OxPhos for ATP production in male mice under injury-induced conditions. In alveolar macrophages, UCP2 has been reported to limit a pathogen-killing (pro-inflammatory) phenotype while promoting a pre-resolving (anti-inflammatory) phenotype by inducing efferocytosis of pathogen debris (Better et al., 2023). These studies suggest that UCP2 may provide metabolic flexibility, allowing cells to select the most appropriate metabolic substrate in response to different metabolic challenges and polarization states. Given these observations, the variation in metabolic pathways between LPS-MΦ and IL4-MΦs suggests that UCP2 levels may be regulated during macrophage polarization.

In this study, we differentiated bone marrow-derived macrophages into pro-inflammatory and anti-inflammatory subsets and exposed them to various metabolic challenges, such as shortage or deprivation of glucose, glutamine and pyruvate, as well as hypoxia-mimicking conditions. We analyzed UCP2 protein levels alongside oxygen consumption rate (OCR), extracellular acidification rate (ECAR), cell proliferation, and oxygenation. We aimed to evaluate whether changes in environmental parameters influence adaptive shifts in UCP2 levels and to determine if these changes correlate with the distinct metabolic profiles of pro- and anti-inflammatory macrophages under both physiological and pathological conditions.

## 2. Materials and Methods

### 2.1 Isolation of tissue-resident macrophages

8–12-week-old C57BL/6J mice were euthanized by cervical dislocation, and tissues (spleen, lung, liver, bone marrow, brain, colon, adipose tissue, peritoneum) were rapidly collected. The tissues were minced into small pieces and digested using the Multi Tissue Digestion Kit (130-110-203, Miltenyi Company, California, USA) for 20 minutes. After digestion, single cell suspensions were filtered through a 70 µM strainer and centrifuged at 500 g for 3-5 minutes. The pellet was then resuspended in EasySep buffer (20144, StemCell, Cologne, Germany) and passed through a 40 µm strainer into 5 mL tubes. Rat serum was added to the cell suspensions, followed by incubation with the first antibody (APC or PE conjugated; #100-0033, EasySep™ Release Mouse APC/PE Positive Selection Kit, StemCell, Cologne, Germany) for 5 minutes at room temperature. Magnetic beads were then added, and the mixture was incubated for 3 minutes at room temperature. The cells were topped up with EasySep buffer and placed into the EasyEights™ EasySep™ Magnet (#18103, StemCell, Cologne, Germany) for 3 minutes. After incubation, the supernatant was carefully aspirated without touching the cells adhering to the side facing the magnet. The tube was then removed from the magnet, and cells were washed with Easy Sep buffer before being returned to the magnet for another 3-minute incubation. This wash step was repeated twice. After the final wash, cells were resuspended in EasySep buffer, and release buffer concentrate was added, followed by a 3-minute incubation. Subsequently, tubes were placed back in the magnet, and the supernatant containing the positively selected cells was collected. The cells were centrifuged, resuspended in EasySep buffer, and the entire procedure was repeated with a second antibody (PE or APC conjugated, using a different fluorophore than in the first step). After the second enrichment step, cells were resuspended in Triazole and total RNA was isolated by using Monarch RNA Cleanup Kit (T2040, New England Biolab Company, Massachusetts, USA) according to the manufacturer’s instructions.

### 2.2 Isolation and culture of murine BMDMΦs and RAW 264.7 cells

Eight to ten weeks old female C57Bl/6j mice were purchased from Janvier Labs (Le Genest-St-Isle, France). Mice were sacrificed by cervical dislocation in accordance with the ethical approval of the Austrian national authority under the Animal Experiments Act (Tierversuchsgesetz 2012; approval number ETK-172/11/2023). The femora, tibiae, and humeri were then harvested, and bone marrow was collected by rinsing the bones with media using a syringe.

Cells were seeded at Nunc™ EasYDish™ Dishes (150468, Thermo Fisher Scientific Inc., Massachusetts, USA) and cultured under standard conditions (37°C, 5% CO_2_, and 95% humidity) for 5 days in RPMI 1640 medium (2522621, Gibco, Dublin, Ireland), supplemented with heat-inactivated fetal bovine serum (HI FBS; 10082147, Gibco, Dublin, Ireland), 1% penicillin/streptomycin (P/S) (15140122, Gibco, Dublin, Ireland), and with 25 ng/mL macrophage colony-stimulating factor (M-CSF; 315-02-50UG, Peprotech, Gibco, Dublin, Ireland). Media change and re-addition of MCSF was done at day 3 of culture.

On day 5, bone marrow-derived macrophages (BMDMΦs) were harvested by scraping, counted, and reseeded into appropriate cell plates for overnight culture with the same media. For immunoblotting and quantitative PCR analysis, 1.5 × 10^6^ cells and for proliferation assay, 25 × 10^5^ cells were seeded per well in 6-well plates (174901, Thermo Fisher Scientific Inc., Massachusetts, USA). For Seahorse extracellular analysis, 80 × 10^4^ cells were seeded per well in Seahorse 96XFe plates (102959-100, Agilent, California, USA).

After overnight culture, macrophages were exposed to DMEM media without glucose, glutamine, or phenol red (A1443001, Gibco), but supplemented with 10% HI FBS, 1% P/S and with 15 ng/mL M-CSF. Unless otherwise specified, the media also contained 2 mM GlutaMAX (35050061, Gibco), 5.5 mM glucose (A2494001, Gibco), and 1 mM sodium pyruvate (11360070, Gibco).

To induce an inflammatory phenotype in BMDMΦs, 50 ng/mL of lipopolysaccharide (LPS; L8643, Sigma-Aldrich, Massachusetts, USA) and 5 ng/mL of interferon-γ (INF-γ; 315-05, Peprotech, Gibco) were added to the media. To polarize BMDMΦs into an anti-inflammatory phenotype, 20 ng/mL of interleukine-4 (IL-4; 214-14, Peprotech, Gibco) and 20 ng/mL of interleukin-13 (IL-13; 200-13, Peprotech, Gibco) were added. If required for the experiment, 25 mM of lactate (1614308, Sigma Aldrich, Massachusetts, USA), 10 µM MG132 (M8699, Sigma-Aldrich), and 10 or 50 µM of UK5099 (5.04817, Sigma Aldrich) were added.

RAW 264.7 macrophage-like cells, kindly provided by Dr. Reinhard Gruber (Oral Biology group at the Dentistry school of Vienna), were expanded in RPMI 1640 medium supplemented with 10% FBS, 1% P/S for 4 passages. Cells were then seeded at 1 × 10^6^ cells/well for immunoblotting and 25 × 10^5^ cells/well for proliferation assays in 6-well plates. Polarization of RAW 264.7 cells was performed following the same protocol used for BMDMΦs.

### 2.3 Protein isolation and quantitative immunoblot analysis

Protein isolation from cells and immunoblot blot analysis were performed as described in Rupprecht et al. (Rupprecht et al., 2014). In brief, cells were washed with PBS, collected in RIPA-buffer containing a 1:50 protease inhibitor cocktail, and sonicated. Total cellular protein was isolated after centrifugation and quantified using a BCA kit (A55860, Thermo Fisher Scientific Inc., Massachusetts, USA). We loaded 20 µg of total cellular protein per lane for immunoblot analyses. We used the following primary antibodies: SDHA (ab14715, Cambridge, United Kingdom), OGDH (15212-1-AP, Proteintech Group Inc, UK), VDAC (ab14734, Abcam Inc., UK), β-actin (A5441, Sigma Aldrich Inc., Massachusetts, USA), and self-designed, validated antibody against UCP2 (Rupprecht et al., 2012; Rupprecht et al., 2014). Antibodies against STAT1 (9172, Cell Signaling Technology, Inc., Massachusetts, USA), p-STAT1 (9167, Cell Signaling Technology, Inc., Massachusetts, USA), STAT6 (9162, Cell Signaling Technology, Inc., Massachusetts, USA), p-STAT6 (9361, Cell Signaling Technology, Inc., Massachusetts, USA), bcl2 (3498, Cell Signaling Technology, Inc., Massachusetts, USA), BAX (2772, Cell Signaling Technology, Inc., Massachusetts, USA), NFKBIA/IkB alpha (H-4) (sc-1643, Santa Cruz Biotechnology Inc., USA), and α-tubulin (801202, Bio Legend Inc., San Diego, USA) were used at dilutions 1:1000, and 1:10000, respectively. Immunoreactions were detected by luminescence using a secondary antibody against rabbit (7074, Cell Signaling Technology, Inc., Massachusetts, USA) or mouse (NA931V, GE HealthCare Technologies, Inc., Chicago, USA). Antibodies were linked to the horseradish peroxidase and ECL Immunoblot Blotting reagent (1705061, Bio-Rad Laboratories Ges.m.b.H., California, USA). The intensity of bands was quantified using Software VisionWorks 8.20 (Analytik Jena GmbH+Co., Jena, Germany). For all Western blot analyses in this study, band intensity normalization (Fig. S1G) was performed as follows: (1) For each blot, UCP2 and SDHA intensity values for each macrophage subset (UCP2_(MΦ)_, SDHA_(MΦ)_) were divided by the corresponding values from spleen tissue (UCP2_(Sp)_, SDHA_(Sp)_), resulting in UCP2_(MΦ)_/UCP2_(Sp)_ and SDHA_(MΦ)_/SDHA_(Sp)_. (2) The relative protein amount (I_rel_) was calculated as I_rel_ = ITP/IHK, where ITP is the intensity of the target protein and IHK is the intensity of the normalization control (SDHA, β-actin, or α-tubulin). Therefore, UCP2_(MΦ)_/UCP2_(Sp)_ values were divided by SDHA_(MΦ)_/SDHA_(Sp)_ values to correct for mitochondrial loading. (3) UCP2/SDHA ratios for LPS-MΦ and IL4-MΦ samples were normalized to the ratio of resting MΦ samples.

### 2.4 RNA isolation and quantitative PCR analysis

Gene expression analysis was performed as previously described (Sternberg et al., 2023). In brief, total RNA was isolated from BMDMΦs using TRI-Reagent (RT111, Molecular Research Center, Ohio, USA) following the manufacturer’s instructions. cDNA synthesis was performed using the High-Capacity cDNA Reverse Transcription Kit (AM1722, Applied Biosystems, California, USA), according to the manufacturer’s instructions. For gene expression analysis with qRT-PCR, primers were designed to span exon-exon junctions using NCBI Primer-BLAST. PCR product sizes were kept in the range of 80-150 bp to ensure high primer efficiency. Only primers with amplification efficiencies between 90-105 %, verified through serial dilution of cDNA, were used. Amplicon sizes were validated on polyacrylamide gels. Quantitative reverse transcription PCR (qRT-PCR) was performed on a qTower384 real-time PCR system (Analytik Jena GmbH+Co., Jena, Germany) using Luna master mix (M3003, New England Biolab Company, Massachusetts, USA). Plates were loaded in triplicates using 1:1 diluted cDNA and amplification was performed at 62℃ annealing temperature. Murine ribosomal protein L4 (Rpl4) was used as a housekeeping gene. CT values for *Ucp2*, hypoxia-inducible factor 1α gene (HIF1A), Egl Nine homolog 1 gene (*Egln1*), and *Rpl24* were determined for each sample. Primer sequences are listed in Supplementary Table 1. Ucp2 mRNA expression was normalized to mRpl4 levels.

### 2.5 Determination of oxygen consumption rate (OCR) and extracellular acidification rate (ECAR)

BMDMΦs were seeded in Seahorse 96XFe plates at a density of 80,000 cells per well and incubated overnight in RPMI medium supplemented with 2 mM glutamine, 10 mM glucose, 1 mM pyruvate, 10% FBS, and 1% P/S. For nutrient starvation experiments, the entire plate was washed with PBS and refilled with fresh media (control or nutrient starved). Polarization was performed for four and 18 hours as previously described.

One hour before the experiment, the media were replaced with Seahorse XF RPMI assay medium (103681, Agilent, California, USA), supplemented with the specified concentrations of glucose (103577, Agilent, California, USA), glutamine (103279, Agilent, California, USA), and pyruvate (103578, Agilent, California, USA). OCR and ECAR were measured in parallel using an XFe96 extracellular flux analyzer (Agilent, California, USA). The compounds for the Seahorse XF Cell Mito Stress Test -4.5 µM oligomycin (75351, Sigma Aldrich, Massachusetts, USA), 4.5 µM FCCP (C2920, Sigma Aldrich, Massachusetts, USA), 2.5 µM rotenone (557368, Sigma Aldrich, Massachusetts, USA), and 1.25 µM antimycin A (A8674, Sigma Aldrich, Massachusetts, USA) - were added according to the manufactureŕs instructions. Each independent experiment was performed on a new plate. The minimal number of wells per condition for each experiment was five for basal measurements and three for the Mito Stress Test and mitochondrial nutrient usage inhibition analysis. OCR and ECAR values for each well were normalized to the total protein concentration of the cells seeded in that well.

### 2.6 Proliferation assay

BMDMΦs or RAW 264.7 cells were seeded in 6-well plates at a density 25 × 10^5^ cells per well and incubated overnight in control media. For nutrient shortage experiments, the entire plate was washed with PBS and refilled with fresh media (control or nutrient-deprived), followed by polarization as described previously. Cell proliferation was monitored using the Incucyte^®^ S3 Live-Cell Analysis System (Sartorius Lab Instruments GmbH & Co., Göttingen, Germany). This real-time live-cell analysis assay allows continuous monitoring and quantification of cell growth over time. The software was programmed to take 16 pictures every 2 hours over a period of one to two days.

### 2.7 RNA sequencing data processing

Raw reads were analyzed with a Nextflow v23.10.1 workflow, which included the following steps: analysis of read quality, alignment of the reads, and transcript quantification (Di Tommaso et al., 2017). The reads quality was verified by FastQC v0.12.1 with default parameters, followed by index building and read alignment against the GRCm39 reference genome with STAR v2.7.11, and using the Gencode release M34 GTF file as the annotation parameter (Dobin et al., 2013) (https://www.bioinformatics.babraham.ac.uk/projects/fastqc). The following parameters were specified for STAR alignment: --outSAMtype BAM SortedByCoordinate –outSAMunmapped Within. Next, an indexed transcriptome was built with Gencode Mouse Release M34, followed by quantification of transcripts. Indexing and quantification were performed using Salmon v1.10.3 0.3 (Patro et al., 2017) with default parameters for index building. For quantification, the following parameters were specified: --gcBias --validateMappings. All reference datasets were obtained from Gencode (https://www.gencodegenes.org/mouse). MultiQC was used to aggregate quality reports (Ewels et al., 2016)

### 2.8 Unsupervised clustering of the TRM gene count matrices

K-Means clustering was performed with R stats package v4.3.1 (Chambers et al.). The evaluation of clustering quality was assessed using the fviz_nbclust function from the R factoextra package v1.0.7 (Kassambara, 2020), with the following methods: elbow method, silhouette method and gap statistic. To examine gene expression differences between samples for Ucp2, a CPM gene activity matrix was used. Gene-specific changes based on Z-score values (calculated as the difference between a specific sample’s gene expression level and the mean divided by the standard deviation) were visualized with pheatmap package (Pheatmap: pretty heatmaps version 1.0.12 from CRAN)(Kolde, 2019).

### 2.9 Statistics

Gene and protein expression data were analyzed using GraphPad Prism software 7 (GraphPad Software, San Diego, CA, USA). Relative mRNA expression was calculated using the 2^-ΔΔCt^ method. Statistical analysis of the data was performed using one-way analysis of variance (ANOVA), with the Kruskal-Wallis test for non-parametric data and Dunn’s multiple corrections. Protein expression was analyzed with one-way ANOVA using Dunnett’s multiple comparisons test. A significance level was set to 0.05. Significant differences are indicated as follows: *P < 0.05, **P < 0.01, ***P < 0.001.

## 3. Results

### 3.1 BMDMФs differentiated from monocytes express UCP2

Monocytes were differentiated into macrophages using macrophage colony-stimulating factor (MCSF) and then polarized into pro-inflammatory (LPS-MΦ) or anti-inflammatory (IL4-MΦ) macrophages using LPS/INFγ and IL4/IL13, respectively (Fig. 1A). The success of the differentiation was verified by the presence of phosphorylated STAT1 in LPS-MΦ and STAT6 in IL4_MF after 18 hours (Murray et al., 2014) (Fig. 1B).

**Figure 1.**
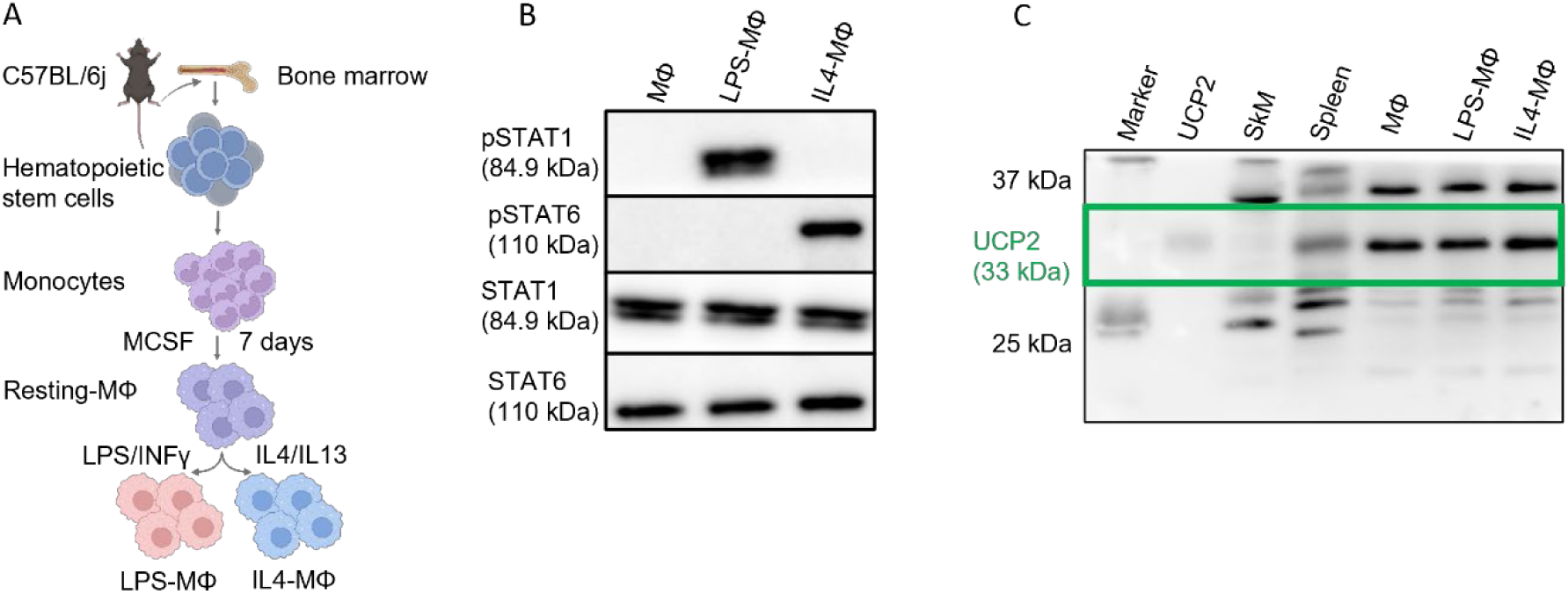
UCP2 abundance in polarized bone marrow-derived macrophages. (A) Scheme of differentiation and polarization of bone marrow-derived macrophages into pro- and anti-inflammatory phenotypes. (B) Representative WB showing STAT1 and STAT6 as well as their phosphorylated forms (pSTAT1 and pSTAT6) in LPS-MΦ and IL4-MΦ, respectively. (C) Representative WB confirming the expression of UCP2 in pro-ad anti-inflammatory macrophages. Recombinant mouse UCP2 (1 ng) and spleen, and skeletal muscle (SkM) were used as positive and negative controls for UCP2 expression, respectively. 20 µg total cell or tissue protein was loaded per lane.

Immunoblot analysis confirmed the presence of UCP2 protein in all BMDMΦ subsets similar to monocytes (Fig. 1C). To avoid the challenges associated with nonspecific commercial antibodies in UCP2 research, which have led to controversial results in various studies, we used a custom-designed polyclonal anti-UCP2 antibody (Rupprecht et al., 2012) that has been previously validated and applied in independent studies (Beikbaghban et al., 2024; Maes et al., 2023). The specificity of the detected bands was confirmed by positive signals in the applied positive controls (recombinant mouse UCP2, mouse spleen) and their absence in the negative control (skeletal muscle, SkM) (Fig. 1C).

### 3.2 UCP2 protein levels correlate with oxygen consumption rates in anti-inflammatory macrophages

To evaluate whether UCP2 protein levels vary upon macrophage polarization under physiological macronutrient conditions (5.5 mM glucose, 2 mM glutamine, and 1 mM pyruvate), we performed immunoblot analysis of BMDMΦs at different time points of polarization. The results showed a decrease in UCP2 levels in LPS-MΦ compared to IL4-MΦ (Fig. 2A).

**Figure 2.**
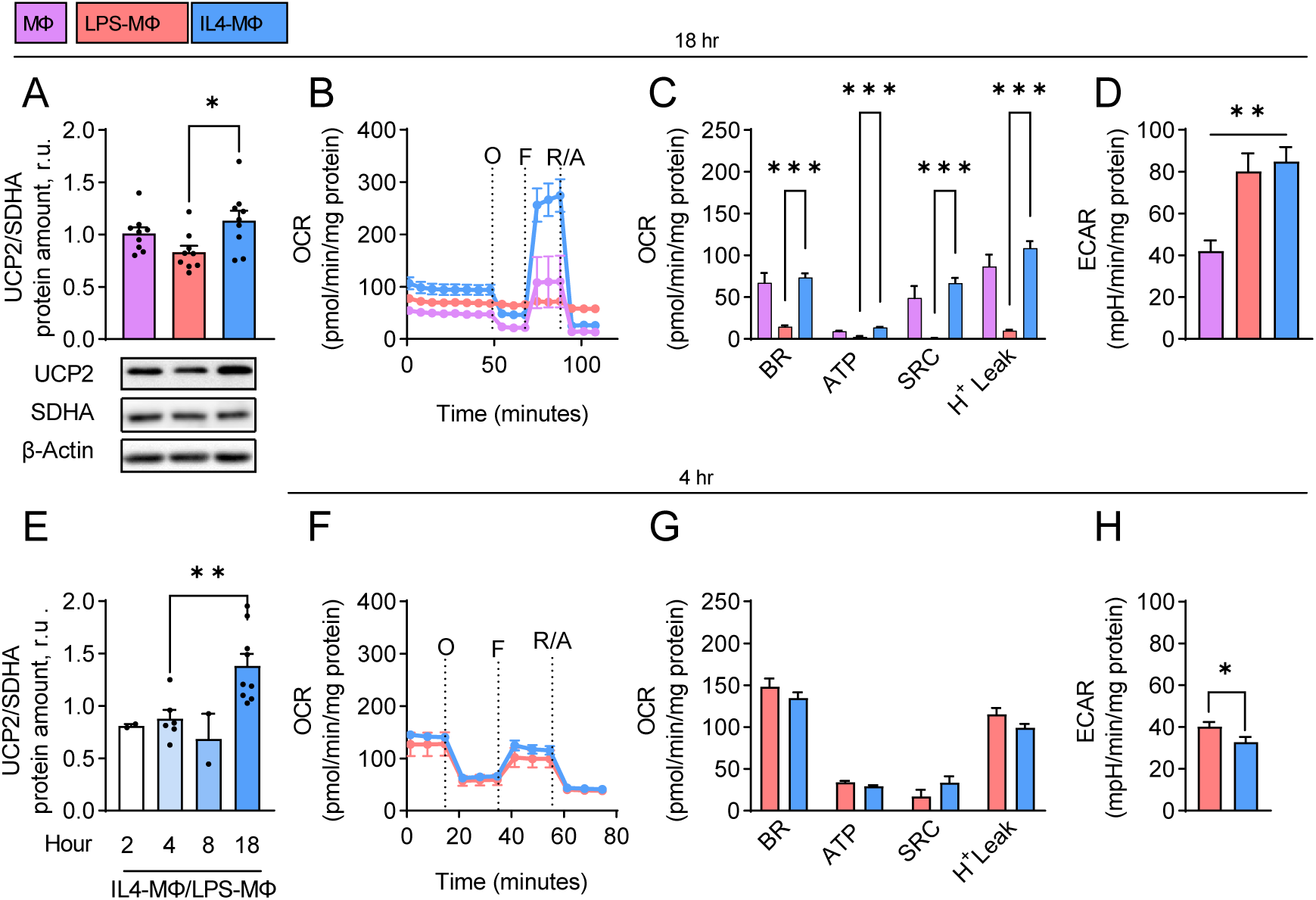
Correlation of polarization state and metabolism of BMDMΦs with UCP2 protein level under physiological nutrition. (A) Representative WB of UCP2, SDHA, and β-actin with quantification analysis of UCP2/SDHA (N=9), (B) representative OCR, (C) quantification of OCR-derived parameters and (D) ECAR (N=5) in MΦs, LPS-MΦs and IL4-MΦs after 18 hours of polarization under physiological macronutrient conditions. (E) Ratio of quantification analysis of UCP2/SDHA in IL4-MΦs to LPS-MΦs following 2, 4, 8, and 18 hours of polarization under physiological macronutrient condition, (F) representative OCR, (G) quantification of OCR-derived parameters, and (H) ECAR (n=4) in MΦs, LPS-MΦs and IL4-MΦs after 4 hours polarization under physiological macronutrient condition. 20 µg of isolated total protein from each group was loaded per lane. Data are presented as mean ± SEM, \**p*< 0.05, \*\**p*< 0.01 \*\*\**p*< 0.001. O, oligomycin; F, FCCP; R/A, rotenone/antimycin. BR, basal respiration; SRC, spare respiratory capacity.

Importantly, the ratio of succinate dehydrogenase (SDHA) to actin, used as a control for mitochondria amount, remained unchanged (Fig. S1A). Oxygen consumption rate (OCR) analysis showed a flat curve for LPS-MΦ (Fig. 2B), consistent with the decreased basal respiration of LPS-MΦ compared to IL4-MΦ (Fig. 2C).

However, acidification as an indirect indication of glycolysis-produced lactate amount, measured by extracellular acidification rate (ECAR) was similar between the two macrophage subsets (Fig. 2D). In general, IL4-MΦs showed increased mitochondrial activity, as spare respiratory capacity, mitochondria-related ATP production and proton leak were much higher than in LPS-MΦs (Fig. 2C).

To explore the dynamics of UCP2 protein regulation during polarization, we conducted a time-course experiment. Notably, during the first 8 hours of polarization, UCP2 protein levels in IL4-MΦ and LPS-MΦ did not differ significantly. However, after 18 hours, UCP2 levels were significantly higher in IL4-MΦ vs. LPS-MΦ (Fig. 2E) under physiological macronutrient levels.

Both LPS-MΦ and IL4-MΦ responded similarly to respiratory chain inhibitors, showing respiring mitochondria as early as 4 hours post-polarization (Fig. 2F). Basal respiration, ATP production, and proton leak were comparable between the two phenotypes (Fig. 2G). Initial signs of glycolytic metabolic adaptions were visible as LPS-MΦ showed a higher ECAR than IL4-MF after 4 hours of polarization (Fig. 2H). In line with literature (Pecqueur et al., 2001), *Ucp2* mRNA and protein levels were differentially altered upon 18 hours of polarization, as analysis of non-polarized and polarized BMDMΦs under physiological macronutrient conditions showed no regulation of *Ucp2* mRNA levels relative to *mRpl4* (Fig. S2A).

Furthermore, the proliferation rate of non-polarized and polarized BMDMs was assessed over 48 hours using an Incucyte scanning device. The results showed no increase in proliferation rates in any phenotype of BMDMΦs (Fig. S2D), suggesting that macrophages do not proliferate once they are polarized into the inflammatory and anti-inflammatory subsets. However, this result is only partially supported by other studies, as some studies on anti-inflammatory macrophages have shown that their proliferation is regulated by P53 (Li et al., 2015; Motta et al., 2021).

### 3.3 UCP2 levels remain elevated in anti-inflammatory macrophages in the absence of glucose or glutamine

To investigate the impact of glucose deprivation on UCP2 expression in macrophages, we performed immunoblot analysis on polarized macrophages under glucose-free conditions. The results showed a decrease in UCP2 levels in LPS-MΦ compared to IL4-MΦ after 18 hours of polarization in the absence of glucose, although no significant difference was observed after 4 hours (Fig. 3A).

**Figure 3.**
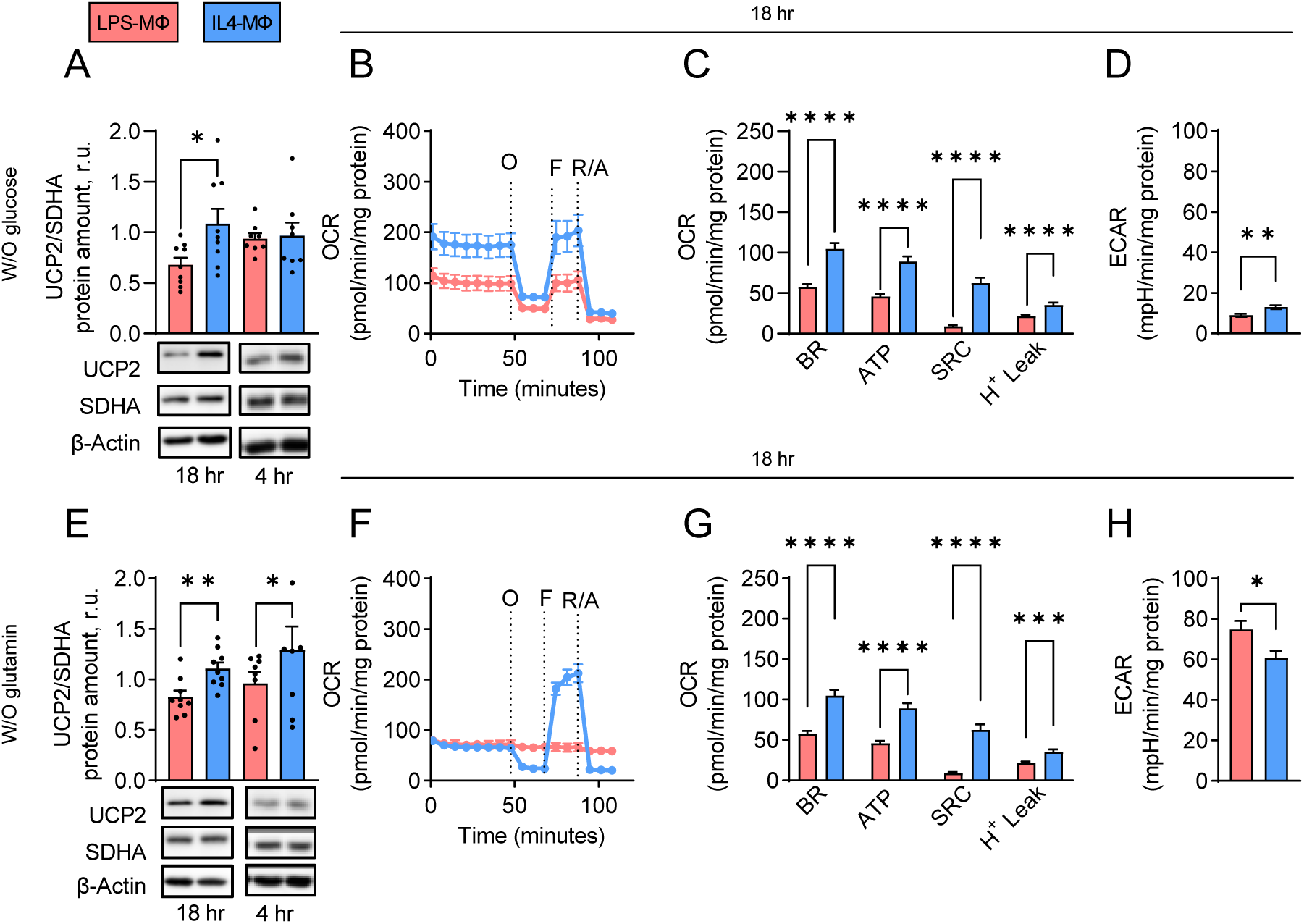
Correlation of polarization state and metabolism of BMDMΦs with UCP2 protein level in the absence of glucose. (A-D) or glutamine (E-H). (A) Representative WB of UCP2, SDHA, and β-actin with quantification analysis of UCP2/SDHA (N=9), (B) representative OCR, (C) quantification of OCR-derived parameters and (D) ECAR (N=5) in LPS-MΦs and IL4-MΦs after 18 or 4 hours polarization in the absence of glucose. (E) Representative WB of UCP2, SDHA, and actin with quantification analysis of UCP2/SDHA (N=9), (F) representative OCR, (G) quantification of OCR-derived parameters and (H) ECAR (N=5) in LPS-MΦs and IL4-MΦs after 18 or 4 hours polarization in the absence of glutamine. 20 µg of isolated total protein from each group was loaded per lane. Data are presented as mean ± SEM, *p< 0.05, **p< 0.01, ****p< 0.0001. O, oligomycin; F, FCCP; R/A, rotenone/antimycin. BR, basal respiration; SRC, spare respiratory capacity.

In the absence of glucose, mitochondria from LPS-MΦ remained respiring and responded to inhibitors of respiratory complexes (Fig. 3B). However, these respirating mitochondria exhibited a lower level of all OCR parameters in comparison to IL4-MΦs in the absence of glucose (Fig. 3C). Notably, the amount of UCP2 protein was lower in LPS-MΦs lacking glucose in their media (Fig. 3 A) compared to the levels of UCP2 in the presence of glucose in the media (Fig 2A). This finding suggests a regulatory effect of glucose on the level of UCP2 in LPS-MΦs.

Likewise, ECAR was lower in LPS-MΦ than in IL4-MΦ (Fig. 3D). ECAR was especially low due to the absence of glucose, which results in lower lactate production. After 4 hours of polarization, both basal respiration and other respiratory parameters, including ATP production, respiratory capacity, and proton leak were similar between both phenotypes (Fig. S3A). Notably, LPS-MΦ exhibited lower ECAR than IL4-MΦ after 4 hours (Fig. S3B), consistent with the results from the overnight incubation.

Immunoblot analysis of macrophages polarized in the absence of glutamine showed reduced amounts of UCP2 in LPS-MΦ compared to IL4-MΦ after both 18 and 4-hour incubations (Fig. 3E). As observed under physiological nutrition and in the presence of glucose but absence of glutamine mitochondria of LPS-MΦ were not respiring, resulting in a flat OCR (Fig. 3F). To investigate whether lactate, a major byproduct of glucose metabolism in LPS-MΦs (O’Neill and Pearce, 2016) could account for the lack of respiratory chain (RC) activity in these cells (as previously suggested in alveolar epithelial type II cells (Lottes et al., 2015)), OCR analysis was performed on LPS-MΦ polarized for 18 hours in the absence of glucose. The injection of 25 mM of lactate, followed by RC inhibitors, showed that mitochondria in LPS-MΦ remained respiring, similar to control macrophages (Fig. S3C), excluding lactate as explanation for the observed effects.

Basal respiration and all other respiratory parameters, including ATP production, respiratory capacity, and proton leak, were significantly lower in LPS-MΦ compared to IL4-MΦ after both 18 hours (Fig. 3G) and 4 hours (Fig. S3D) in the absence of glutamine. ECAR was higher in LPS-MΦ than in IL4-MΦ after both 18 hours (Fig. 3H) and 4 hours (Fig. S3E).

RT-PCR analysis of non-polarized and polarized BMDMΦs cultured in the absence of glucose or glutamine for 18 hours showed no difference in UCP2 mRNA levels (Fig. S2B and S2C). This suggests that UCP2 modification occurs at the post-translational level. Neither IL4-MΦ nor LPS-MΦ exhibited any proliferative activity under these conditions (Fig. S2E and S2F). However, gene expression analysis of *Ucp2* in BMDMs did not correlate with UCP2 protein levels under different nutritional conditions or polarization states. Again, this discrepancy aligns with previous studies UCP2 under glucose deprivation conditions (Rupprecht et al., 2019), rotenone-induced oxidative stress (Giardina et al., 2008) or LPS injection (Pecqueur et al., 2001) in pulmonary tissues. Notably, this phenomenon is often attributed to translational and post-translational regulation of UCP2 (Mailloux et al., 2012; Pecqueur et al., 2001; Stanzione et al., 2022). Additionally, UCP2 has a short half-life of less than 30 minutes (Rousset et al., 2007) and is not degraded by the cytosolic proteasome (Azzu and Brand, 2010). However, LPS has been shown to induce proteasomal degradation of UCP2 (Russell and Tisdale, 2009). To address these discrepancies and to test whether MG132, an inhibitor of proteasome degradation, could rescue the reduction in UCP2 protein levels in LPS-MФs, we exposed them to MG132 for two hours before 18 hours polarization. The reduced degradation of IκBα in the presence of MG132 indicates that proteasomal activity is partially, though not completely, inhibited (Fig. S2G). Despite this partial inhibition, we observed no increase in UCP2 levels in the MG132-treated group compared to the untreated control (Fig. S2G). However, because the inhibition was incomplete, we cannot rule out the possibility that proteasomal degradation is responsible for the reduced UCP2 levels observed in LPS-MΦs.

As additional control for our experiments, we polarized the immortalized macrophage-like cell line RAW264.7 into inflammatory and anti-inflammatory phenotypes under different concentrations of glucose, glutamine and pyruvate. In contrast to primary BMDMΦs, UCP2 levels in RAW264.7 cells remained unchanged across all phenotypes. However, regardless of polarization state, RAW264.7 cells were sensitive to glutamine deprivation and expressed significantly lower UCP2 levels in the absence of glutamine (Fig. S4A-G). LPS-RAW264.7 cells exhibited a reduced proliferation rate compared to IL4-RAW264.7 cells under physiological macronutrient conditions and after 12 hours of glucose deprivation (Fig. S4 H-J). This difference can be attributed to the proliferative, immortalized nature of RAW264.7 cells, in contrast to the non-proliferative behavior of BMDMΦs (Fig. S2D, E, and F).

In summary, these experiments demonstrate that primary macrophages maintain higher UCP2 protein levels in relation to mitochondrial respiration, independent of nutrient availability. In contrast, UCP2 expression in RAW cells is more dependent on glutamine availability, resembling the behavior observed in murine neuroblastoma cells (N18TG2) (Beikbaghban et al., 2024; Rupprecht et al., 2019).

### 3.4 Anti-inflammatory macrophages express higher UCP2 protein levels in the absence of glucose

To investigate whether pathologically high concentrations of glucose, as seen in metabolic syndrome patients, and glutamine modulate UCP2 levels in BMDMs, polarized and non-polarized BMDMΦs were incubated with various concentrations of (i) glucose (0-25 mM) with constant concentration of glutamine (2 mM) (Fig. 4A) and (ii) glutamine (0-4 mM) with constant concentration of glucose (5.5 mM) (Fig. 4B) for 2, 4, and 18 hours.

**Figure 4.**
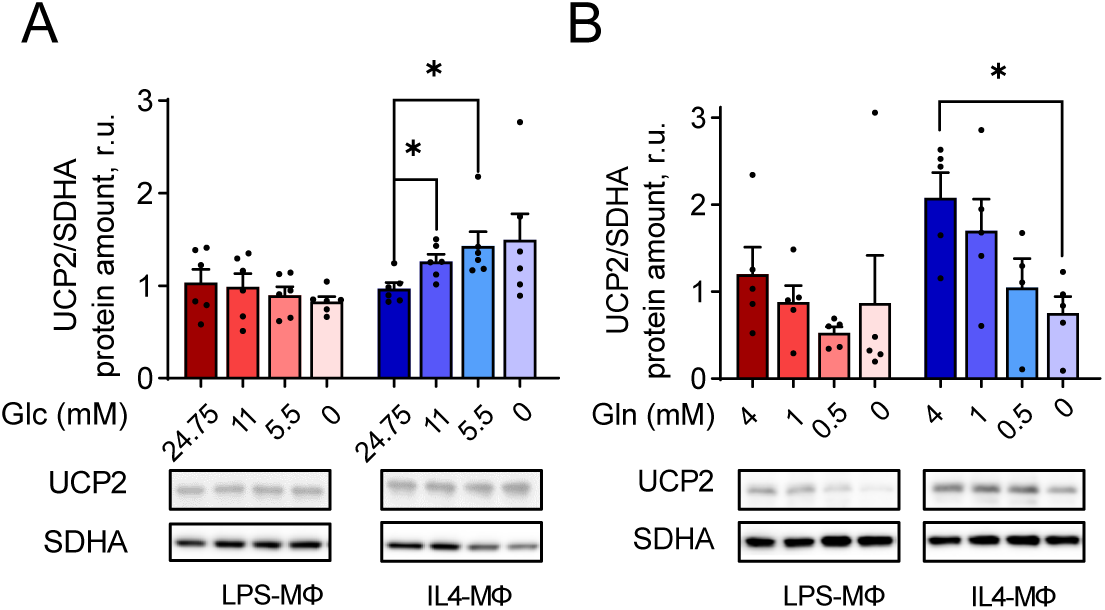
Protein levels of UCP2 under different concentrations of glucose and glutamine. Representative WB and quantification analysis of UCP2/SDHA LPS-MΦs and IL4-MΦs after overnight polarization and incubation under (A) 24.75-, 11.1, 5.5, and 0 mM glucose in the presence of 2 mM glutamine or (B) 4-, 2-, 0.5-, and 0-mM glutamine in the presence of 5.5 mM glucose (N=6). 20 µg of isolated total protein from each group was loaded per lane. Data are presented as mean ± SEM, *p< 0.05.

Immunoblot analysis of BMDMΦs revealed a significant increase in UCP2 levels in IL4-MΦ but not in LPS-MΦ when incubated with 5.5 mM and 11 mM glucose compared to 25 mM glucose after 18 hours of incubation (Fig. 4A). Decreasing the glutamine concentration reduced UCP2 expression in IL4-MΦ after 18 hours of incubation (Fig. 4B). UCP2 protein levels remained unchanged under different doses of glucose and glutamine after 2 and 4 hours (Fig. S5 C-D and S5 G-H). Collectively, these results suggest that anti-inflammatory macrophages predominantly rely on glycolysis rather than mitochondrial respiration when glucose concentrations are higher, thereby reducing their dependence on UCP2’s C4 metabolite transport function and on OxPhos in general.

### 3.5 UCP2 levels decreased in the absence of pyruvate

It has been proposed that in neuroblastoma cells, UCP2 transports C4 metabolites from the mitochondrial matrix to the cytosol during glucose shortage, providing substrates for conversion into pyruvate, which in turn fuels the TCA cycle (Rupprecht et al., 2019). Consistent with this, previous RNA sequencing data from K562 UCP2 knockout cells identified pyruvate kinase R/L (PKRL) as the gene with the highest fold change compared to control K562 cells (Beikbaghban et al., 2024).

Building on this, we examined the impact of pyruvate levels on UCP2 expression across different macrophage subsets. Pyruvate was excluded from the media, resulting in three distinct nutritional conditions: no pyruvate, no glucose/no pyruvate, and no glutamine/no pyruvate. Immunoblot analysis of LPS-MΦ showed a significant reduction in UCP2 levels in the absence of pyruvate (Fig. 5A) and when both pyruvate and glucose were excluded (Fig. 5B). Notably, this effect was dependent on glutamine availability, as UCP2 levels remained unchanged when pyruvate was removed in the absence of glutamine (Fig. 5C). The ratio of SDHA to actin remained unchanged in all experimental conditions (Fig. S6A-C).

**Figure 5.**
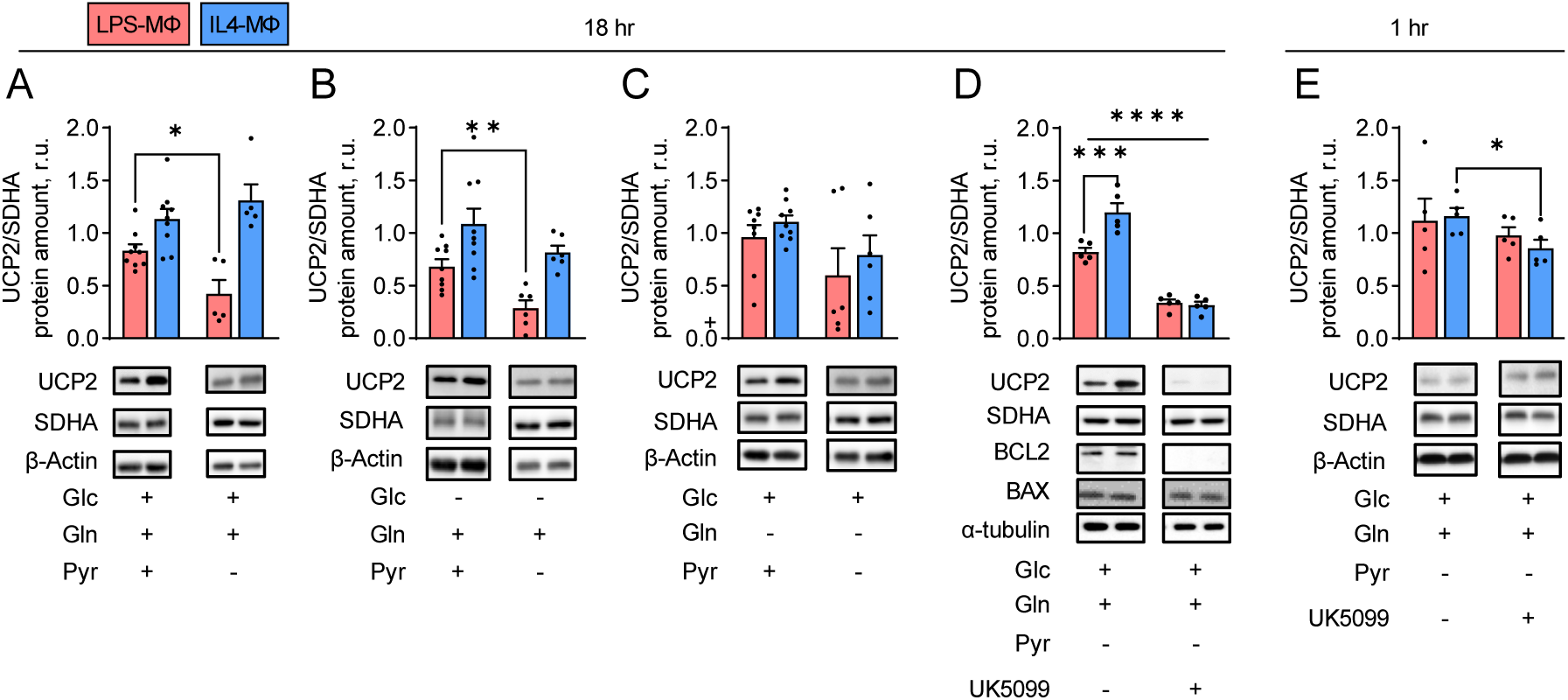
UCP2 levels in the absence of pyruvate or after blocking its uptake into mitochondria. Representative WB of UCP2, SDHA, β-actin, Bcl2, BAX, and α-tubulin and quantification analysis of UCP2/SDHA in LPS-MΦs and IL4-MΦs after 18 hours polarization and incubation under physiological macronutrient conditions vs. absence of pyruvate (A), absence of glucose vs absence of glucose and pyruvate (B), absence of glutamine vs absence of glutamine and pyruvate (N=6-9) (C), and blocking of the mitochondrial pyruvate carrier with UK5099 for overnight (D) and 1 hour (E) (N=5). 20 µg of isolated total protein from each group was loaded per lane. Data are presented as mean ± SEM, *p< 0.05, **p< 0.01, ***p< 0.001, ****p< 0.0001. Glc, glucose; Gln, glutamine; Pyr, pyruvate.

We also investigated whether blocking pyruvate entry into the mitochondria using UK5099 as a selective inhibitor of the mitochondrial pyruvate carrier (MPC) affects UCP2 levels. As shown in Fig. 5D, BMDMΦs exhibited a downregulation of UCP2, while mitochondrial SDHA remained unchanged. The α-tubulin levels decreased after the inhibition of pyruvate insertion, indicating that the cells are under stress due to lack of pyruvate metabolism in the mitochondria. No Bcl2 was detected in macrophages, suggesting that apoptosis was induced after 18 hours of incubation with 10 µM UK5099. We hypothesized that the prolonged treatment may have induced apoptosis, so we reduced the incubation time to 1 hour using 50 µM UK5099. This shorter treatment resulted in a decrease in UCP2 levels in IL4-MΦ compared to control (Fig. 5E). Taken together, these experiments highlight pyruvate as a critical nutrient for mitochondrial respiration and macrophage survival. They also confirm that the reduced UCP2 levels in LPS-MΦ are primarily due to pyruvate being converted to lactate, rather than being transported into the mitochondria, fueling the TCA cycle and thereby driving C4 metabolite levels.

### 3.6 Hypoxia of anti-inflammatory macrophages reduces UCP2 protein levels

Since lactate production is a hallmark of oxygen deprived conditions which can be triggered also by LPS (Blouin et al., 2004), we addressed the question whether we can mimic the observed effects on UCP2 levels in LPS-MΦ, following exposure of the BMDMΦs to a hypoxic condition. Therefore, macrophages were cultured and polarized in a CoCl_2_-induced hypoxia-mimicking environment for 18 hours. Immunoblot analysis revealed that the hypoxia-mimicking environment resulted in a reduction of UCP2 levels in IL4-MΦ compared to normoxic conditions, bringing UCP2 expression to levels comparable to those observed in LPS-MΦ (Fig. 6A). However, no changes in UCP2 levels were observed in LPS-MΦ under hypoxic conditions, showing an already maximal downregulation effect by LPS alone.

**Figure 6.**
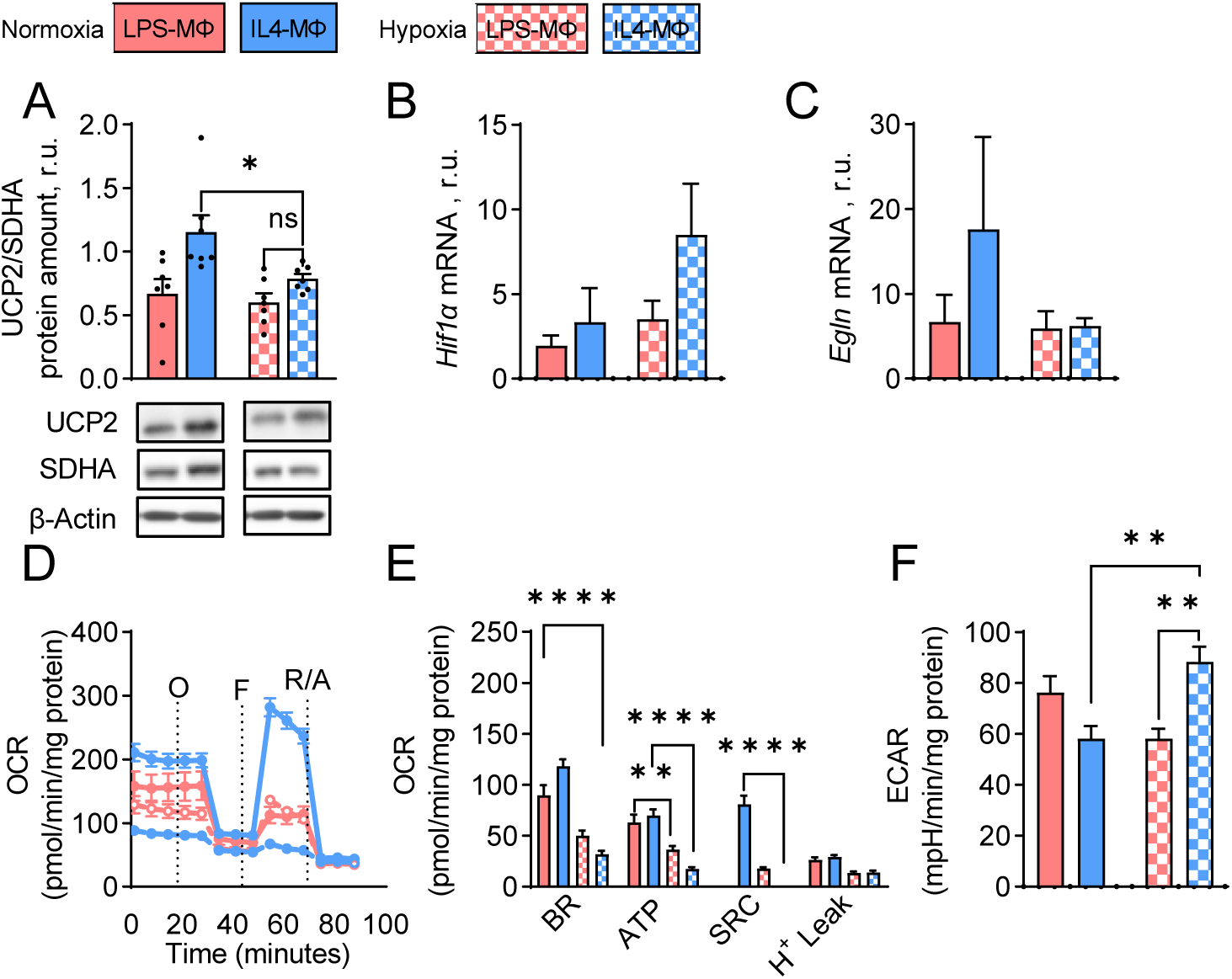
UCP2 levels and metabolism of BMDMΦs under hypoxic-like conditions. (A) Representative WB and quantification analysis of UCP2/SDHA in LPS-MΦs and IL4-MΦs after overnight polarization and incubation under physiological micronutrient conditions in the absence (normoxia) or presence of CoCl2 (hypoxia-like conditions). (N=7). (B) QRT-PCR analysis of (B) Hif1α and (C) Egln1 genes with mitochondrial ribosomal protein l4 (Rpl4) as mitochondrial reference gene in LPS-MΦs and IL4-MΦ (N=4). (D) Representative OCR, (E) quantification of OCR-derived parameters and (F) ECAR (N=4) in LPS-MΦs and IL4-MΦs after overnight polarization in physiological nutrition under normoxia or hypoxia-like conditions. 20 µg of isolated total protein from each group was loaded per lane. Data are presented as mean ± SEM,, *p< 0.05, **p< 0.01, ****p< 0.0001. O, oligomycin; F, FCCP; R/A, rotenone/antimycin. BR, basal respiration; SRC, spare respiratory capacity. Amount of the mRNA is in relative units to Rpl4 as a housekeeping gene and calculated using the 2^-ΔΔCt^ method.

SDHA protein levels remained unchanged under hypoxia (Fig. S7A). IL4-MΦs showed increased *Hif1a* and *Engl1* mRNA under hypoxia mimicking conditions (Fig. 6B and C). Extracellular flux analysis revealed a decrease in basal OCR, ATP production, and spare respiratory capacity in IL4-MΦs under hypoxia compared to normoxia. Conversely, ECAR of IL4-MΦs increased upon exposure to the hypoxia-mimicking environment. These results strengthen the previously postulated mechanism of UCP2’s pyruvate-lactate-oxygen dependency. Additionally, the data suggest that the CoCl2-induced hypoxia-mimicking environment stabilizes HIF-1α, which in turn reduces PHD2, leading to a decreased need for UCP2’s transport activity and, consequently, its abundance.

### 3.7 Variability in UCP2 mRNA levels among tissue-resident macrophages

To place the data in a broader context and highlight the significance of our findings, we analyzed whether Ucp2 expression varies across different murine tissue-resident macrophages (TRMs). RNA sequencing analysis revealed that TRMs from the spleen exhibited the highest Z-score for Ucp2 mRNA (Fig. 7), followed by blood monocytes, bone marrow-derived macrophages (BMDMs), and lung-associated macrophages.

**Figure 7.**
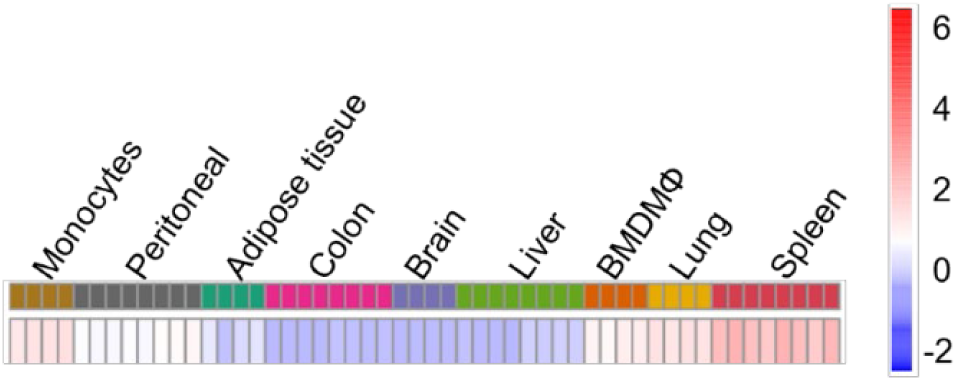
RNA sequencing analysis of tissue-resident macrophages. Heatmap illustration of UCP2 mRNA Z-score in different tissue-resident macrophages (TRMs) resulted from RNA sequencing analysis of TRMs from 4 to 8 different mice. The different colors represent the tissue of origin of the macrophages, with each individual mouse contributing a different column.

Peritoneal macrophages displayed a relatively lower Z-score, indicating a basal level within the grading scale. Interestingly, macrophages derived from adipose tissue, brain, colon, and liver showed negative Z-scores for Ucp2 mRNA (Fig. 7). These results demonstrate significant variation in Ucp2 mRNA expression across different TRM populations. However, since mRNA levels do not always correlate with protein expression, further studies are needed to assess the distribution of UCP2 protein levels among the TRMs.

## 4. Discussion

The present study demonstrates that IL4-polarized macrophages exhibit higher UCP2 levels than LPS-polarized macrophages after 18 hours of polarization (Fig. 8). The increased UCP2 expression in IL4-MΦs was associated with a higher oxygen consumption rate (OCR). Furthermore, reducing glucose concentration from a pathological level to zero, resulted in an increase in UCP2 expression in IL4-MΦs. However, when glutamine concentration was also reduced under glucose-deprived conditions, UCP2 levels decreased. Notably, blocking pyruvate entry into mitochondria for 18 hours led to a loss of UCP2 protein in both macrophage phenotypes. Under hypoxia-mimicking conditions induced by CoCl2, IL4-MΦs showed a significant reduction in UCP2 expression and OCR, which brought UCP2 levels closer to those observed in LPS-MΦs.

**Figure 8.**
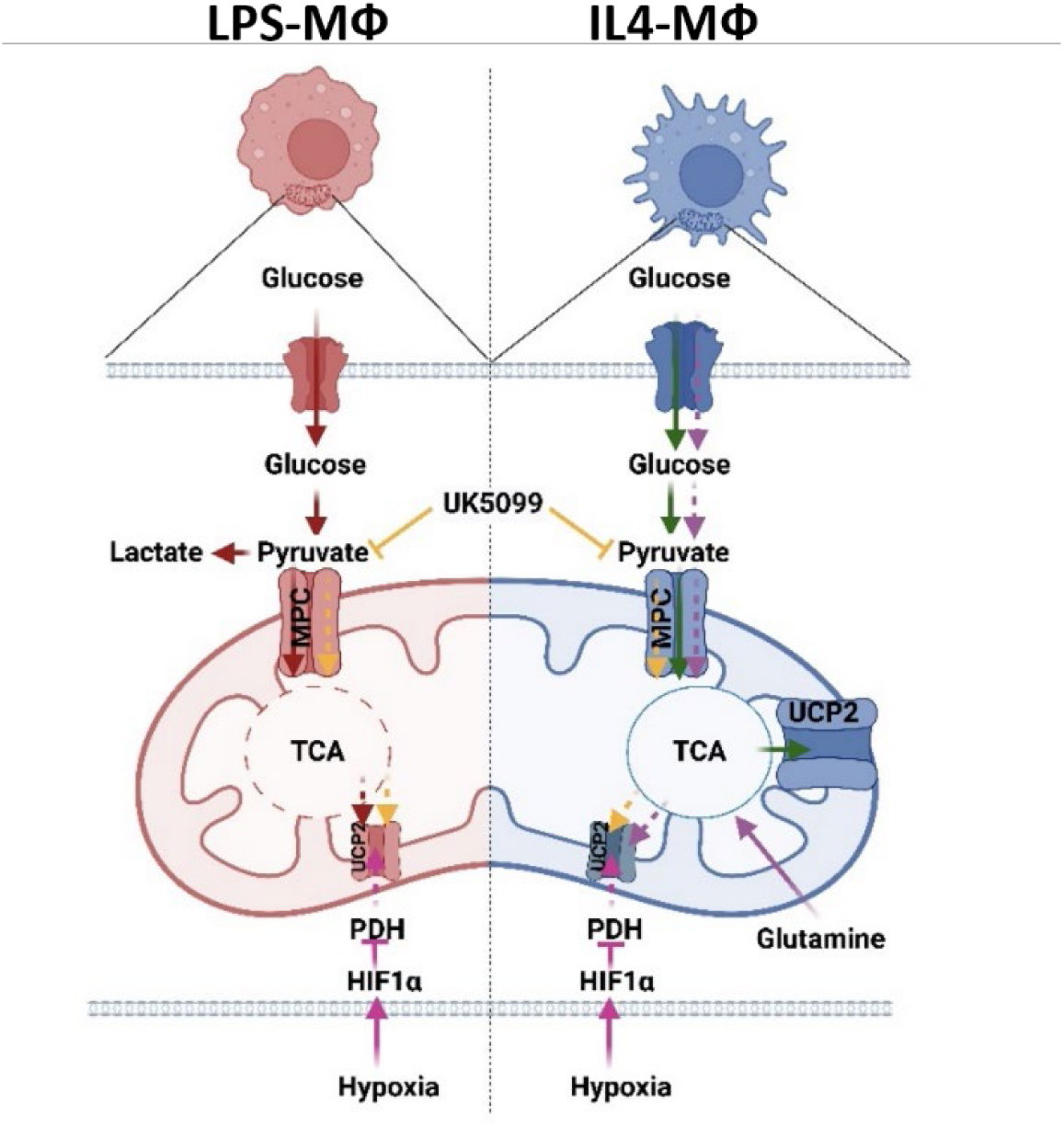
UCP2 regulation based on available metabolic substrate in the microenvironment of macrophages’ subsets. Red arrows: Entry of physiological dose of glucose (1) lead to high level of glycolysis in LPS-MФs ending up to high lactate production and release (2), resulting in low UCP2 protein level (3). Green arrows: In IL4-MФs pyruvate remains as the end product (2), entering the TCA cycle, resulting in high UCP2 protein level (3). Purple arrays: In IL4-MФs, under glucose concentrations which are lower than 25 mM (1), there is a high consumption of glutamine as alternative substrate (2), resulting in a high UCP2 protein amount (3). Orange arrays: Inhibition of MPC with UK5099 (1) blocks entrance of pyruvate into mitochondria (2), causing almost no expression of UCP2 (3) in both subsets of macrophages. Pink arrays: Hypoxic condition results in increased level of HIF1α (1) which leads to inhibition of pyruvate hdehydrogenase (PDH) (2) and finally decreased expression of UCP2 (3).

### 4.1 IL4-MΦs have increased UCP2 levels compared to LPS-MΦ, which correlate with OCR

Our study confirmed the presence of UCP2 in macrophages, consistent with previous studies investigating UCP2 primarily in immune cells such as macrophages (Alves-Guerra et al., 2003; Arsenijevic et al., 2000). We compared UCP2 levels between LPS-MΦs and IL4-MΦ, whereas most other in vitro investigations on UCP2 in BMDMΦs, have focused solely on LPS-MΦs (Emre et al., 2007a; Gupta et al., 2017; Moon et al., 2023; van Dierendonck et al., 2020), without including a comparison to IL4-MΦs (Faas et al., 2021). Our investigations revealed that UCP2 levels were higher in IL4-MΦs than in LPS-MΦs, which was in line with a previous study in human primary macrophages (Lang et al., 2023). It is also worth noting that some studies on UCP2 in macrophages did not use validated antibodies and appropriate positive and negative controls for the UCP2 band in Western blots, which may be the reason to produce contradictory results by different groups. For example, UCP2 levels were observed to decreased in peritoneal macrophages after LPS treatment (Cortez-Pinto et al., 1998; Emre et al., 2007b; Lee et al., 1999) whereas no changes in UCP2 levels were observed in LPS-treated microglial cells after 18 hours (De Simone et al., 2015).

The correlation between UCP2 levels and oxygen consumption rate in IL4-MΦs suggests that UCP2 supports cells with higher mitochondrial respiration by facilitating the transport of metabolic substrates into the TCA cycle. This role appears to be less relevant for cells that primarily rely on glycolytic metabolism. These findings indicate that the intrinsic metabolism of each macrophage phenotype determines UCP2 level as a downstream target, which contrasts with a study suggesting that UCP2 determines the metabolism and thereby phenotype of the macrophages (Lang et al., 2023).

In contrast to our observation of flat OCR in LPS-MΦs after 24h of polarization in the presence of glucose, investigations on primary human macrophages showed normal OCR in LPS-MΦs after 48 h of treatment (Lang et al., 2023). Additionally, we demonstrated that lactate addition does not cause inactivity of the mitochondria in LPS-MΦs in the absence of glucose, which contradicts previous findings that lactate suppresses respiration due to elevated NADH/NAD^+^ levels (Li et al., 2022). The discrepancies between the results may be attributed to the different cell models being used in their studies which were HepG2 cells and CD8T cells (Cai et al., 2023) or overnight treatment of the cells instead of its acute injection in OCR analysis (Wang et al., 2022). However, other studies have shown that nitric oxide (NO), which is produced to a significant extent by LPS-MΦs, is one of the main inhibitors of mitochondrial respiration (Christopoulos et al., 2021; Van den Bossche et al., 2016).

Similar ECAR levels between the pro- and anti-inflammatory phenotypes under physiological macronutrient conditions indicate a high level of glycolysis in the IL4-MΦs after 18 hours. Previous studies have shown increased ECAR in IL4-MΦs after 18 hours, likely due to their ability to repolarize to the classical pro-inflammatory phenotype (Lundahl et al., 2022; Van den Bossche et al., 2016). The higher ECAR in IL4-MΦs compared to LPS-MΦs in the absence of glucose warrants further investigation into its underlying mechanisms. This difference may be attributed to IL4-MΦś greater ability to repolarize to classical pro-inflammatory macrophages in the presence of glutamine but absence of glucose. Conversely, in the absence of glutamine but with glucose, ECAR was higher in LPS-MΦs than IL4-MΦs, as expected. This indicates a high conversion rate of pyruvate-derived glucose to lactate, resulting in a high ECAR. However, the lack of glutamine appears to limit IL4-MΦ’s ability to repolarize to classical macrophages. Further studies are needed to elucidate the role of metabolic substrates such as glucose and glutamine in IL4-MΦ’s capacity to repolarize to the classical macrophages.

### 4.2 UCP2 levels in Il4-MΦs are determined by the availability of metabolic substrates

In this study, we explored for the first time the impact of different metabolic substrates shortages, such as glucose, glutamine, and pyruvate, on BMDMΦs. We found that expression of UCP2 was significantly lower under pathologically high concentrations of glucose compared to lower concentrations. This phenomenon may be attributed to elevated glycolysis and reduced mitochondrial respiration under elevated glucose levels. As a result, the abundance of UCP2, a transporter essential for supporting mitochondrial respiration, is reduced, as it is less critical in cells relying on glycolysis rather than mitochondrial respiration.

Moreover, the level of UCP2 in IL4-MΦs was found to be sensitive to the concentration of glutamine. This finding supports previous observations in cell lines such as N18TG2, HT29, and INS-1, as well as our current results in RAW264.7 cells. These studies collectively suggest that glutamine plays a key role in regulating UCP2 expression (Hurtaud et al., 2007; Rupprecht et al., 2019; Vozza et al., 2014). In contrast, LPS-MΦs maintained the same basic level of UCP2 regardless of glucose and glutamine concentrations. This can be explained by the fact that LPS-MΦs predominantly rely on glycolysis to produce ATP for their antibacterial activity (Nonnenmacher and Hiller, 2018), while exhibiting an impaired TCA cycle (Jha et al., 2015). This makes the phenotype less dependent on the TCA cycle and, consequently, on the regulation of UCP2 protein levels, which is proposed to provide TCA intermediates for active mitochondrial respiration (Rupprecht et al., 2019). Vozza et al.(Vozza et al., 2014) showed that UCP2 limits oxidation of the pyruvate in the absence of glucose in HepG2 cells, resulting in reduced accumulation of TCA intermediates in the cytosol. This effect was attributed to the role of UCP2 in exporting TCA intermediates, such as pyruvate, from the mitochondria to the cytosol, whereby reducing the load of intermediates like oxaloacetate, which is necessary for the oxidation of pyruvate-derived acetyl-CoA. Consistent with this, our observation of decreased UCP2 levels in LPS-MΦs due to the omission of pyruvate from the culture medium can be attributed to the fact that LPS-MΦs predominantly rely on glycolysis and convert a significant portion of pyruvate to lactate, rather than transporting it into the mitochondria (Kelly and O’Neill, 2015). In the absence of exogenous pyruvate in the growth medium, this phenotype has limited pyruvate availability for mitochondrial entry, leading to a reduction in TCA cycle activity and mitochondrial respiration, resulting in decreased UCP2 expression. In contrast, the levels of UCP2 in IL4-MΦs were found to be independent of exogenous pyruvate. This may be attributed to active TCA in IL4-MΦs fueled by alternative sources, such as glutamine or fatty acids, as IL4-MΦs obtain acetyl-CoA from both pyruvate-derived glucose and fatty acid beta-oxidation (Van den Bossche and van der Windt, 2018). Thus, UCP2 is expressed in these macrophages at a level sufficient to facilitate the transport of TCA intermediates out of the mitochondria, regardless of pyruvate availability.

The loss of UCP2 after 18 hours of blocking pyruvate entry into mitochondria may result from significantly lower mitochondrial respiration. This is consistent with a decreased cell survival rate, as indicated by the absence of Blc2 and reduced α-tubulin levels. As a result, UCP2 becomes less of a priority for the cells and is more prone to degradation. This suggests that UCP2 is one of the first proteins deemed non-essential during the onset of apoptosis. However, the levels of SDHA and other mitochondrial proteins, such as voltage dependent anion channel (VDAC), and oxoglutarate dehydrogenase (OGDH) (Fig. S6D), remained constant after incubation with UK5099. The unaltered level of SDHA may indicate that its aggregation acts as an apoptosis sensor, and when inhibited, tumor cells can evade apoptotic death (Grimm, 2013).

### 4.3 Hypoxic conditions suppress mitochondrial respiration, resulting in lower UCP2 expression

The reduction of UCP2 in IL4-MΦs under hypoxia suggests that the generally lower level of UCP2 in LPS-MΦs may be due to LPS-induced hypoxia in this phenotype (Blouin et al., 2004). Reduced oxygen availability leads to lower mitochondrial respiration. Under hypoxic conditions, HIF1-α upregulation can inhibit pyruvate dehydrogenase activity (Golias et al., 2016), as observed by the higher levels of Hif1α and lower levels of Egl1 (PDH) in IL4-MΦs under hypoxia-mimicking conditions (Fig. 6B and C). This results in reduced pyruvate availability for conversion to acetyl-CoA. The lower OCR in IL4-MΦs under hypoxia-mimicking conditions, as shown in Fig. 6D and E, further confirms the reduction in mitochondrial respiration.

Conversely, the higher ECAR in IL4-MΦs compared to LPS-MΦs under hypoxia-mimicking conditions (Fig. 6F) is a result of HIF1-α stability, which activates the transcription of glycolytic genes (Chen et al., 2001; Woods et al., 2022). Referring to the proposed ROS-scavenging function of UCP2, it has been shown that hypoxic conditions reduce ROS production (Sen et al., 2024). This could also explain the lower expression of UCP2 under hypoxia, where less ROS production leads to less ROS scavenging activity by UCP2 and thus less expression of this protein by cells. Therefore, the lower UCP2 levels observed under hypoxia can be correlated with lower ROS production (Sen et al., 2024) and diminished mitochondrial respiration (Fig. 6D).

Overall, our data showed that pro-inflammatory macrophages, which exhibit reduced mitochondrial respiration, are less dependent on UCP2. In contrast, anti-inflammatory macrophages, which rely more on mitochondrial respiration, utilize UCP2 extensively. Pyruvate is a key substrate regulating UCP2 levels, linking glycolysis and mitochondrial respiration. These findings suggest that macrophages dynamically adjust their UCP2 levels to meet their specific metabolic needs.

## AUTHORS’ CONTRIBUTIONS

**Conceptualization:** J.N., E.E.P.; **Methodology:** J.N., F.S., A.V., R.S; TK; **Validation:** J.N., E.E.P.; **Formal analysis:** J.N., R.S.; **Investigation:** J.N., F.S., T.B., A.V.; R.S.; **Resources:** E.E.P., T.W.; TK; TR; **Writing original draft:** J.N., E:E.P.; **Review & Editing:** E.E.P; J.N, F.S., A.V, T.B, R.S., T.W., T.R., T.K.; **Visualization:** J.N.; **Supervision:** F.S., E.E.P., T.W., T.R.; **Project administration**: E.E.P.; **Funding acquisition:** E.E.P, T.W. All authors have approved the final version of the manuscript.

## ETHICS APPROVAL STATEMENT

All animal experiments were approved by the Austrian national authority according to the Animal Experiments Act, Tierversuchsgesetz 2012-TVG 2012 (ETK-172/11/2023).

## DATA AVAILABILITY

All data are available in the main article or the supplementary materials and from the corresponding author upon reasonable request.

## ACKNOWLEDGEMENTS

We thank Sarah Bardakji (University of Veterinary Medicine) and Toth-Sonns Lena (University of Applied Sciences Vienna of WKW) for the excellent technical assistance. Open Access Funding by the University of Veterinary Medicine Vienna.

