## Supplemental Figures for "Pro- and anti-inflammatory macrophages adjust UCP2 protein levels based on their intrinsic metabolism and available metabolites"

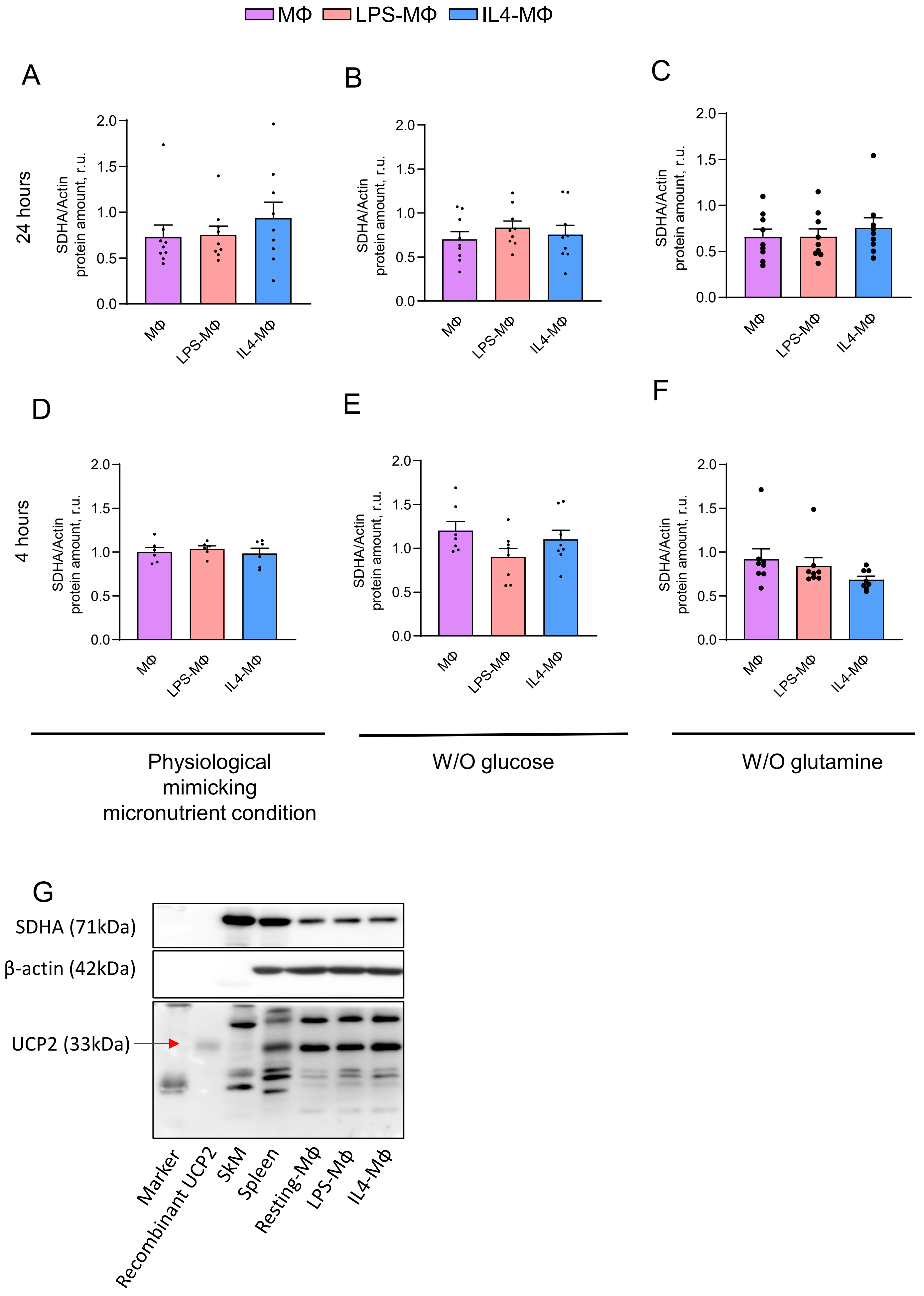


Supplementary Figure 1.

**Supplementary Figure 1. Evaluation of SDHA protein level as mitochondrial protein control in BMDMΦs under different nutritional challenges and normalization of band intensities.** Quantitative analysis for WB of SDHA/Actin in MΦs, LPS-MΦs and IL4-MΦs after 24 hours of polarization in (A) physiological mimicking micronutrient condition, (B) absence of glucose, and (C) absence of glutamine, and 4 hours polarization in (D) physiological mimicking micronutrient level, (E) absence of glucose, and (F) absence of glutamine. Data are presented as mean values $\pm$ SEM, N=6-7. (G) Band intensity normalization was performed in three steps: 1) For each immunoblot, the UCP2 and SDHA intensity values for each macrophage subset (UCP2(MΦ)) were divided by the corresponding UCP2 and SDHA intensity values from spleen (UCP2(Sp)), which were loaded on the same blot, resulting in UCP2(Mφ)/UCP2(Sp) values for each macrophages subset in each blot 2) The UCP2(MΦ)/UCP2(Sp) values were then divided by the SDHA(MΦ)/SDHA(Sp) values. 3) Finally, the UCP2/SDHA ratio for LPS-MΦ and IL4-MΦ samples was normalized against the UCP2/SDHA ratio of resting-MΦ samples.

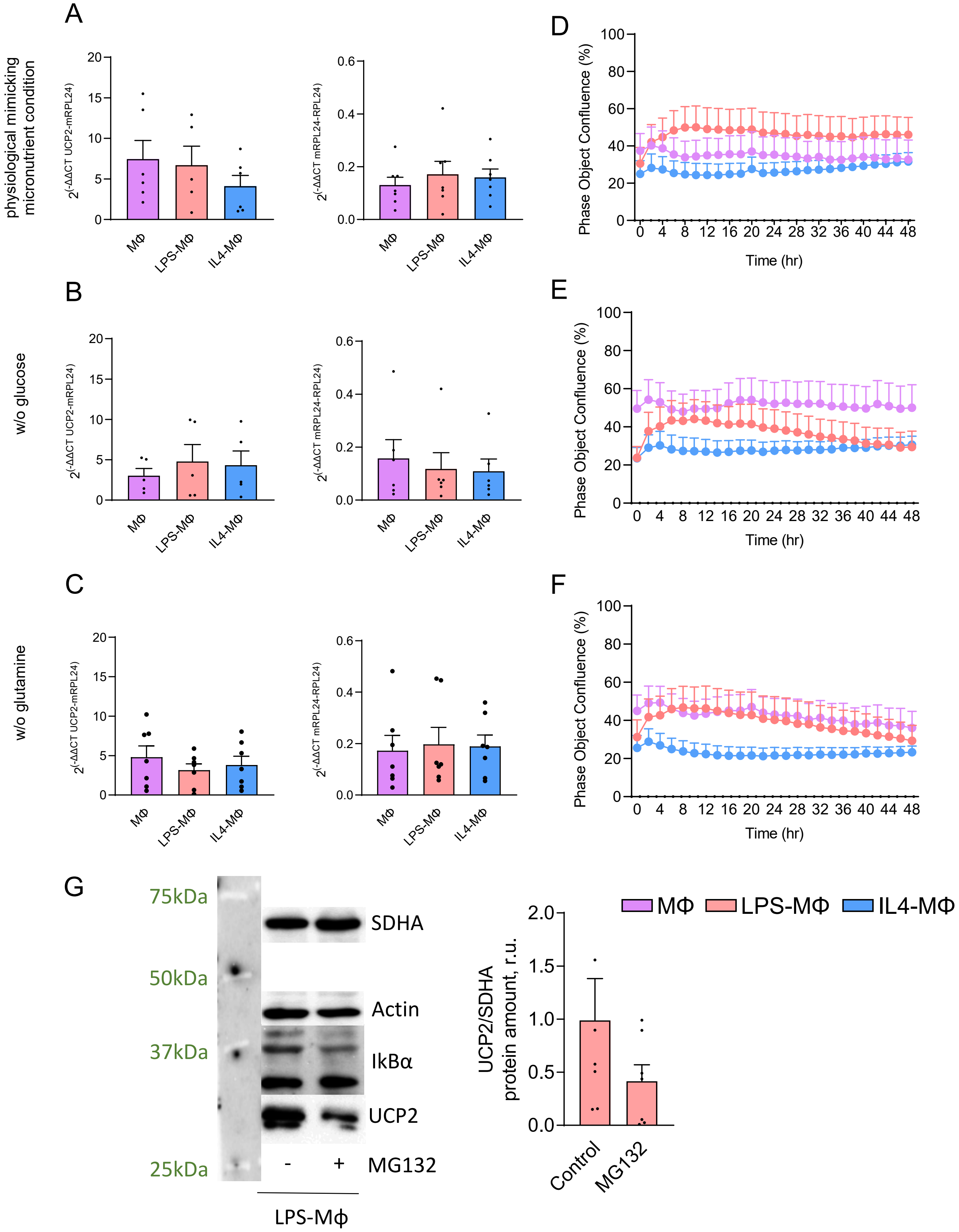


Supplementary Figure 2.

**Supplementary Figure 2. Evaluation of UCP2 gene expression and proliferation of BMDMΦs under different nutritional challenges.** BMDMΦs remained unpolarized (MΦ), or polarized using LPS-MΦ and IL4-Φ in A) physiological mimicking micronutrient condition, (B) absence of glucose, and (C) absence of glutamine overnight followed by total RNA isolation. QRT-PCR analysis of *Ucp2* gene with mitochondrial ribosomal protein L24 (m*Rpl24*) as mitochondrial reference gene and ribosomal protein L24 (*Rpl4*) as cytoplasmic reference gene in MΦs, LPS-MΦs and IL4-MΦ (quantitative analysis of N=5-7 showing mean values ± SEM. BMDMΦs remained either unpolarized (MΦ), or got polarized using LPS+INFγ (LPS-MΦ) and IL4+IL13 (IL4-MΦ) and were grown for 48 h in (D) physiological-mimicking micronutrient level, (E) glucose shortage, and (F) glutamine shortage. Percentage of the cell confluency were measured based on 2 h interval scanning of the cells using Incucyte® SX5 Live-Cell analysis instrument. Quantitative analysis of N=5 showing mean values ± SEM. (G) Representative WB of UCP2, actin, IKBα, and SDHA and quantification analysis of UCP2/SDHA in LPS-MΦs after overnight polarization in the absence or presence of 5μM MG132. 20 μg of isolated total protein of each group is loaded per lane. Data are presented as mean values ± SEM.

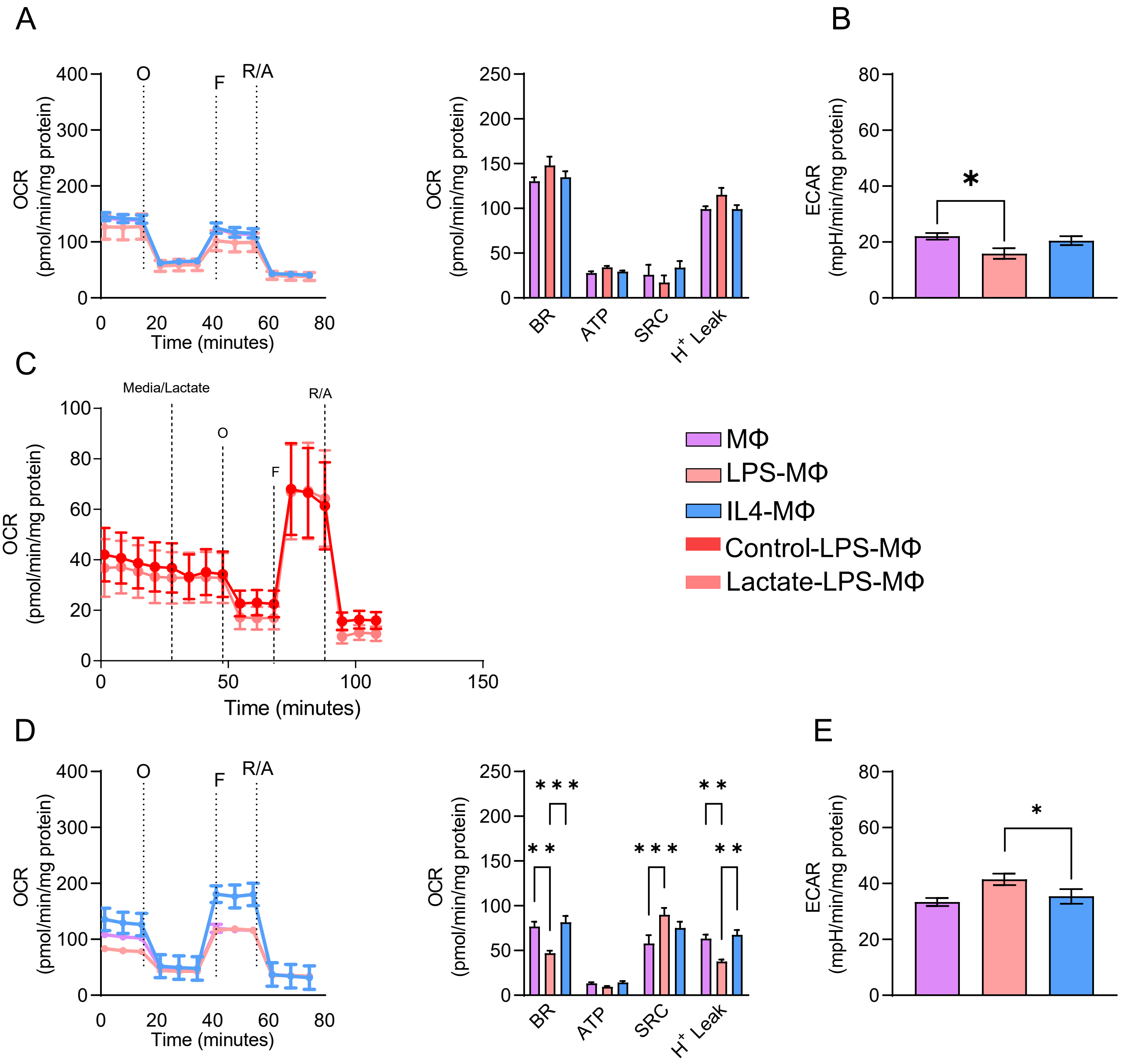


**Supplementary Figure 3. Oxygen consumption rate and extracellular acidification rate of BMDMs in the absence of glucose and glutamine** (A, D) Representative OCR and quantification of OCR-derived parameters and (B,E) ECAR (N=4) in MΦs, LPS-MΦs and IL4-MΦs after four hours polarization in the absence of glucose and glutamine, respectively. (C) OCR of LPS-MΦs in the absence of glucose with and without lactate injection (N=4). Data are presented as mean values ± SEM, *$p < 0.05$, **$p < 0.01$, ****$p < 0.0001$. O, Oligomycin; F, FCCP; R/A, Rotenone/Antimycin.

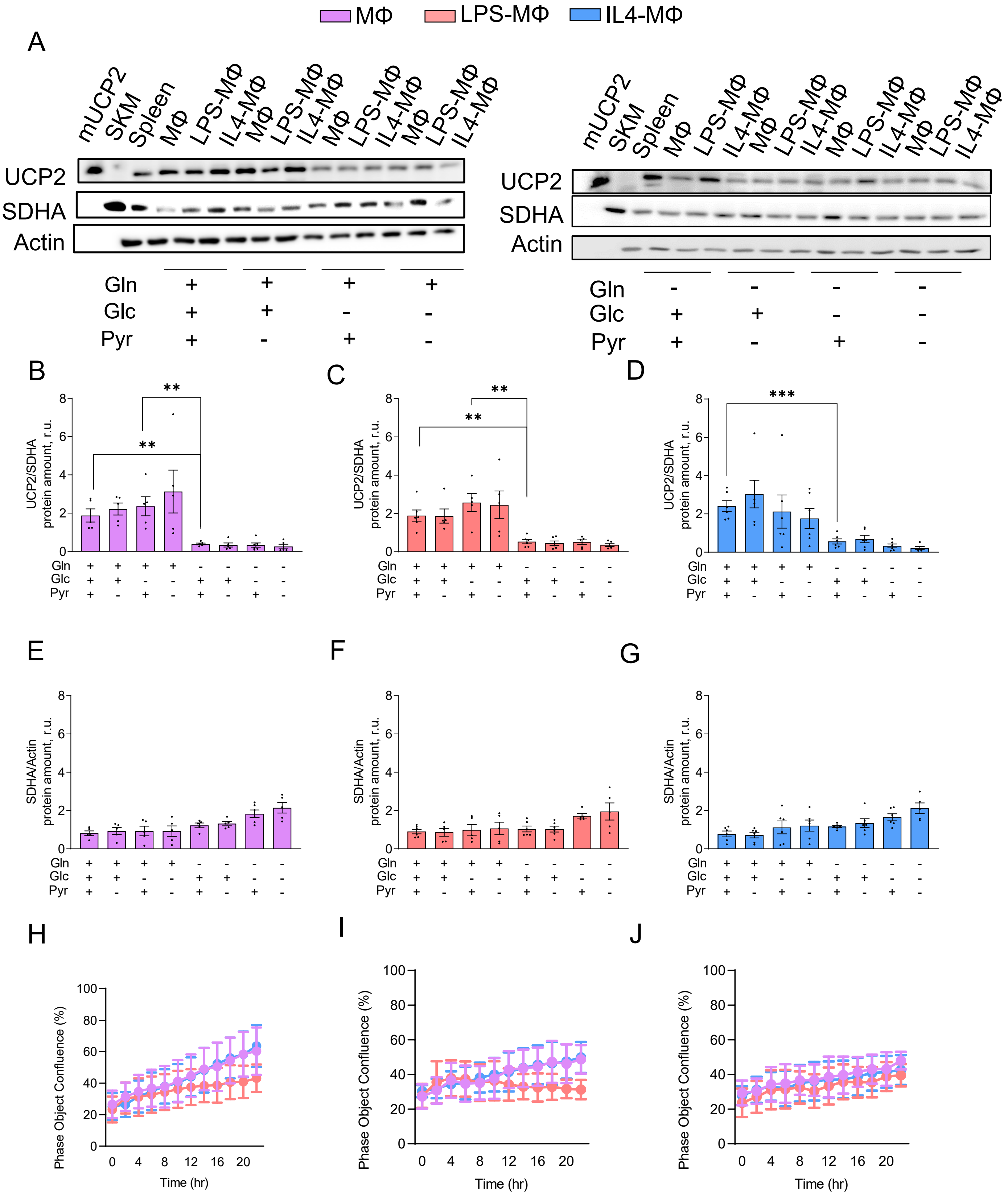


Supplementary Figure 4

**Supplementary Figure 4. Correlation of polarization state of RAW264.7 cells with UCP2 protein level and their proliferation rate.** RAW 264.7 cells remained unpolarized (MΦ) or polarized using LPS/INFɣ and IL4/IL13. Meanwhile, cells were incubated with eight different media containing or not 2mM glutamine, 5.5 mM glucose, and 2mM pyruvate for 24 hr. (A) Representative immunoblots of total cellular protein isolated from the cells using indicated antibodies. (B) Quantitative analysis of five independent experiments of UCP2/SDHA of (B) MΦ, (C) LPS-MΦ, (D) IL4-MΦ, and SDHA/actin of (E) MΦ, (F) LPS-MΦ, (G) IL4-MΦ . Proliferation rate of RAW264.7 cells was evaluated for 24 hr in (H) physiological mimicking macronutrient condition, (I) no glucose, and (J) no glutamine. Percentage of the cell confluency was measured based on 2 hr interval scanning of the cells using Incucyte® SX5 Live-Cell Analysis instrument. Quantitative analysis of N=3 showing mean values ± SEM. Pyr, pyruvate; Glc, glucose; Gln, glutamine.

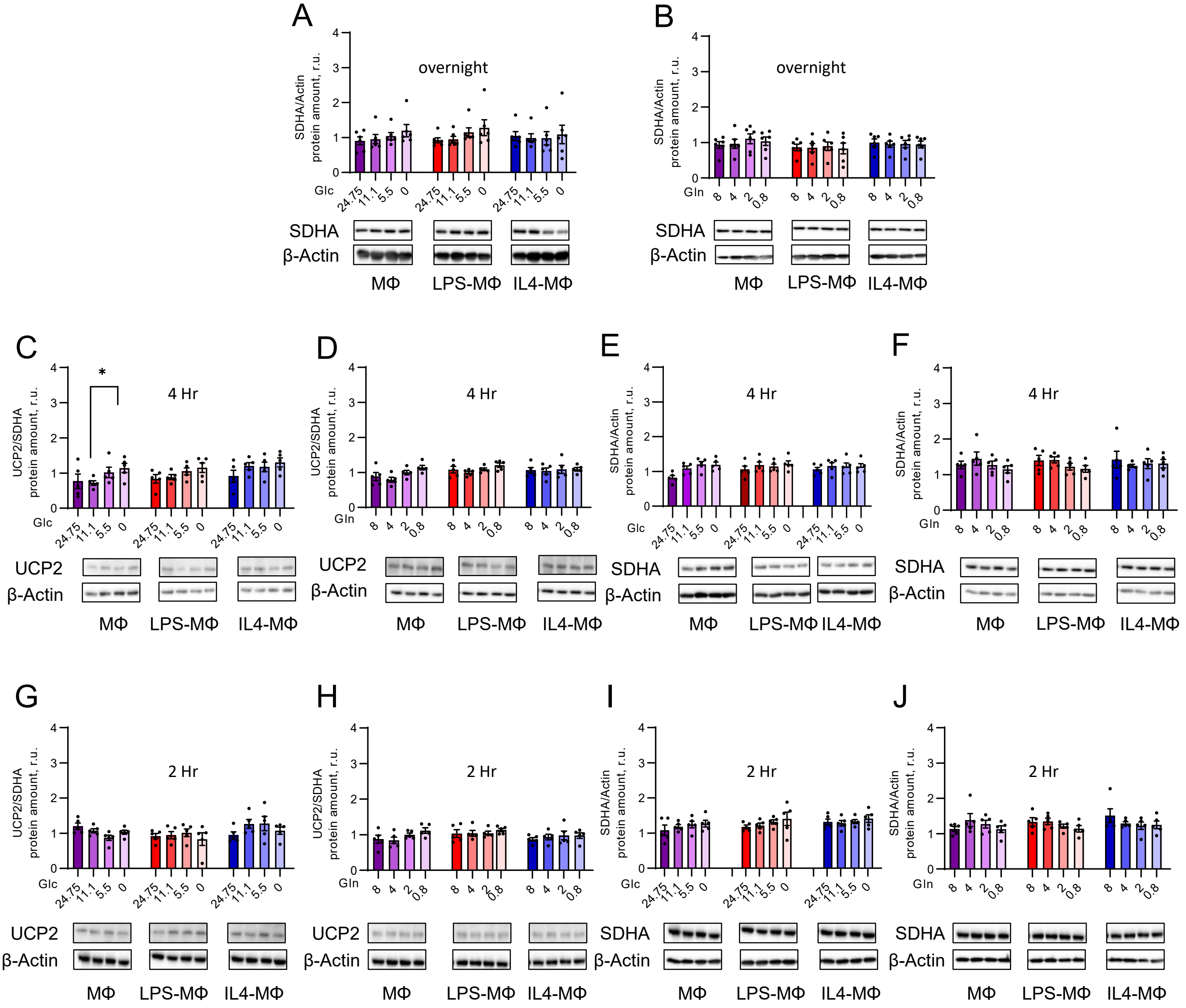


**Supplementary Figure 5. Expression pattern of UCP2 and SDHA as mitochondrial control protein under different doses of glucose and glutamine.** Representative WB and quantification analysis of SDHA/actin in MΦs, LPS-MΦs and IL4-MΦs after (A) overnight (N=6), (E) four hours, (I) two hours (N=5) polarization and incubation in 24.75, 11.1, 5.5, and 0 mM of glucose and 2 mM of glutamine for, Representative WB and quantification analysis of SDHA/Actin in none polarized-Mφs, LPS-MΦs and IL4-MΦs after (B) overnight (N=5), (F) four hours, and (J) two hours (N=5) polarization under 4, 2, 0.5, and 0 mM of glutamine and 5.5 mM of glucose (N=5). Representative WB and quantification analysis of UCP2/actin in none polarized-MΦs, LPS-MΦs and IL4-MΦs under 24.75, 11.1, 5.5, and 0 mM of glucose and 2 mM of glutamine after (C) four hours and (G) two hours (N=5) and 4, 2, 0.5, and 0 mM of glutamine and 5.5 mM of glucose (N=5) after (D) four hours and (H) two hours (N=5). 20 μg of isolated total protein of each group is loaded per lane. Data are presented as mean values ± SEM.

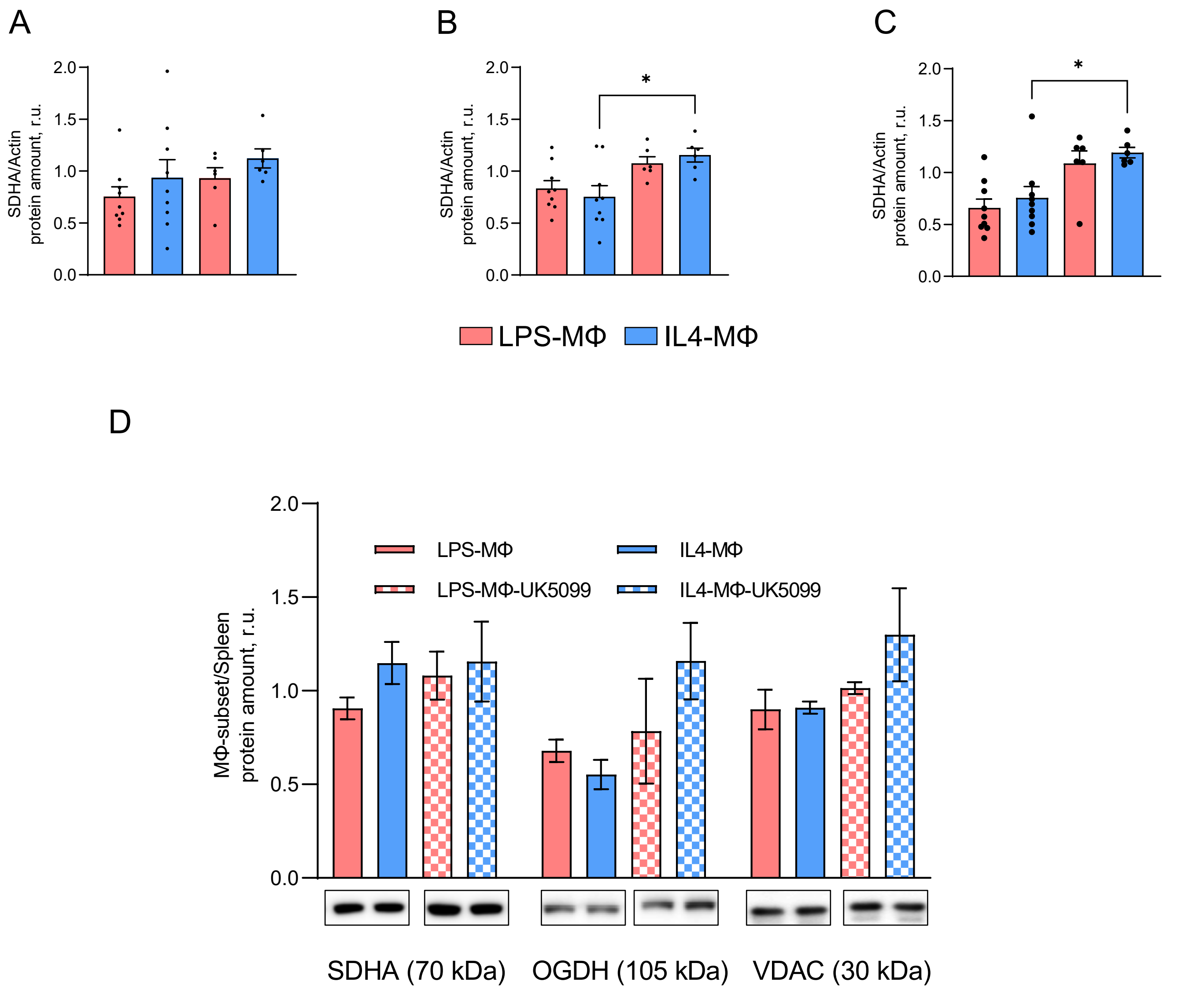


**Supplementary Figure 6. Comparison of SDHA expression as mitochondrial control in the presence versus absence of pyruvate.** Representative WB and quantification analysis of SDHA/Actin in LPS-MΦs and IL4-MΦs after overnight polarization and incubation in the (A)physiological nutrition vs absence of pyruvate, (B) absence of glucose vs absence of glucose and pyruvate, (C) absence of glutamine vs absence of glutamine and pyruvate (N=6-9). (D) Representative WB and quantification analysis of SDHA in each in LPS-MΦs and IL4-MΦs in ratio to spleen after overnight polarization and incubation in the absence and presence of 5μM UK-5099. 20 μg of isolated total protein of each group is loaded per lane. Data are presented as mean values ± SEM, *$p < 0.05$.

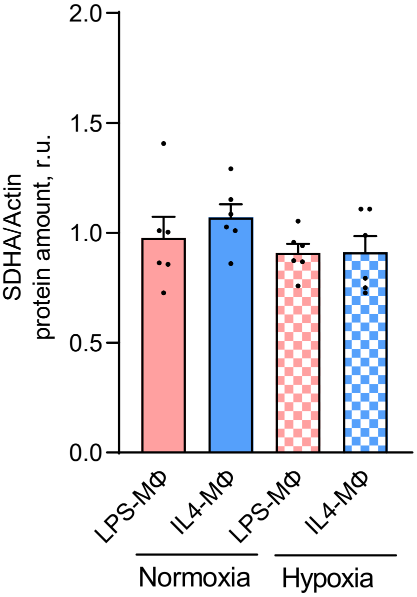


**Supplementary Figure 7. Evaluation of SDHA expression under hypoxia.** Representative WB and quantification analysis of SDHA/actin in LPS-MΦs and IL4-MΦs after overnight polarization and incubation in the physiological nutrition under normoxic versus hypoxic condition (N=7). 20 μg of isolated total protein of each group is loaded per lane. Data are presented as mean values ± SEM, *$p < 0.05$.

**Supplementary Table 1. Primer sequences used for qRT-PCR.**

| Gene | Primer Sequence |
|---|---|
| *Ucp2* | F: 5'-AAAGCAGCCTCCAGAACTCCG-3' |
| | R: 5'-TTCACAGTGGCTGTTGGGGG-3' |
| *Rpl4* | F: 5'-GTATGGCACTTGGCGGAAGG-3' |
| | R: 5'-TGCTCGGAGGGCTCTTTGG-3' |
| *Rpl24* | F: 5'-TGAGCCGTCCAGGTTCCATA-3' |
| | R: 5'-ACAGAATAGGTGCCAGTCTTCA-3' |
| *Hif1a* | F: 5'-AGG ATG AGT TCT GAA CGT CGA AAA G-3' |
| | R: 5'-CAC TGT CTA GAC CAC CGG CA-3' |
| *Egln1* | F: 5'-AAT TCG GCA CGA GGG CAA GT-3’ |
| | R:5'-CAG TGG CGG ATC AGG TCG TC-3' |
